# *Streptococcus* abundance and oral site tropism in humans and non-human primates reflects host and lifestyle differences

**DOI:** 10.1101/2024.05.19.594849

**Authors:** Irina M. Velsko, Christina Warinner

## Abstract

The genus *Streptococcus* is highly diverse and a core member of the primate oral microbiome. *Streptococcus* species are grouped into at least eight phylogenetically-supported clades, five of which are found almost exclusively in the oral cavity. We explored the dominant *Streptococcus* phylogenetic clades in samples from multiple oral sites and from ancient and modern-day humans and non-human primates and found that clade dominance is conserved across human oral sites, with most species falling in the Sanguinis or Mitis clades. However, minor differences in the presence and abundance of individual species within each clade differentiated human lifestyles, with loss of *S. sinensis* appearing to correlate with toothbrushing. Of the non-human primates, only baboons show clade abundance patterns similar to humans, suggesting that a habitat and diet similar to that of early humans may favor the growth of Sanguinis and Mitis clade species.

## Introduction

*Streptococcus* is a diverse and heavily-studied bacterial genus, with a wide range of hosts and habitats including humans and other mammals, amphibians, fish, and food fermentation cultures. While species of this genus include pathogenic host-generalists that infect multiple host species^1^, as well as pathogenic host-specialists^2^, many *Streptococcus* species are commensal host-associated microbiome members that do not inherently cause disease. Phylogenetic analysis of the genus revealed eight well-supported clades, five of which (Sanguinis, Mitis, Anginosus, Salivarius, and Mutans) make up the so-called viridans group^3^. The species within these clades are particularly prominent within the human oral microbiome, exhibiting a highly specific host-niche adaptation within this genus.

In humans, the resident oral *Streptococcus* species exhibit oral site tropism, where particular species preferentially reside on distinct oral surfaces such as the tongue, buccal mucosa, or the tooth surface in dental plaque biofilm^4^. The functional characteristics that distinguish *Streptococcus* species living in different oral niches have been explored in healthy North American populations^4^, which advanced our understanding of the genetic and biochemical drivers of site tropism. However, whether the species partitioning we observe in these populations are characteristic of human populations globally, or whether they are affected by global market integration and urbanization, as well as the evolutionary origins of site-tropism, have not yet been investigated.

*Streptococcus* species play a major role in colonizing tooth surfaces and initiating dental plaque biofilm formation^5^. Recently, *Streptococcus* was shown to be a core genus of the primate dental biofilm by analyzing dental calculus^6^, a mineralized version of dental plaque that forms *in situ* on tooth surfaces during life, and which preserves well in the archaeological record^7^. Fellows Yates, et al.^6^ investigated the distribution of *Streptococcus* in ancient and modern human and non-human primate dental calculus by grouping the *Streptococcus* species by the phylogenetic clades in which they fall, and comparing the abundance of each clade across host species. The streptococcal profiles of ancient humans, including several from Neanderthals, were largely indistinguishable from those of modern humans, yet chimpanzees, gorillas, and howler monkeys each exhibited a distinctive *Streptococcus* clade profile.

In ancient and present-day humans, who have relatively high proportions of *Streptococcus*, the Sanguinis clade was the most abundant clade, while in chimpanzees, who have low proportions of *Streptococcus*, the Anginosus clade was the most abundant^6^. Curiously, however, in a small number of ancient human samples (∼10%), the Sanguinis clade species were nearly absent, and these instead had predominantly Anginosus clade species, strongly resembling the chimpanzee *Streptococcus* clade profiles. Due to the high heterogeneity of samples in that study, no explanation for the chimpanzee-like profile in human samples could be proposed.

Here we investigated the distribution of *Streptococcus* clades in a large dataset of living and ancient human and non-human primate oral samples, to better understand the extent of oral *Streptococcus* site tropism through time and at a global ecological scale. We find that the distribution of *Streptococcus* clades in ancient human calculus is consistent across time and space, with a majority of humans having a Sanguinis and Mitis dominated streptococcal profile, and a minority of humans (∼10%) mostly lacking these clades. Streptococcal profiles are moreover largely consistent within individuals, with teeth across the dentition generally exhibiting similar streptococcal clade patterns. In living populations, the distribution of clades in each oral site is largely consistent across varying levels of global market integration and urbanization, suggesting that diet and lifestyle are not major drivers of streptococcal clade colonization, although species-level differences are observed within clades. Among these is an apparent reduction of *S. sinensis* in the dental plaque of industrialized populations, which may be due to toothbrushing. Each non-human primate investigated has a distinctive *Streptococcus* clade profile, with baboons being most similar to humans. No distinct functional differences underlying site and host specialization were found, suggesting that further characterization of the genes in commensal *Streptococcus* may be necessary to understand niche specialization.

## Results

### Presence of Sanguinis clade species determines clade abundance in ancient humans

We first determined which *Streptococcus* species are found across a large ancient dental calculus dataset comprised of samples from across the globe, spanning 100,000 years, and processed and sequenced in various labs. This allowed us to investigate whether the trend reported by Fellows Yates, et al.^6^, wherein a majority of human ancient dental calculus samples have predominantly Sanguinis clade species and a minority have a chimpanzee-like Anginosus clade-dominated profile, is a universal feature of human ancient dental calculus, or whether this was a feature of the dataset originally used. Following taxonomic profiling with the Genome Taxonomy Database (GTDB), we assigned each *Streptococcus* species in the species table to one of the following previously described *Streptococcus* clades: Sanguinis, Mitis, Salivarius, Anginosus, Bovis, Pyogenic, Mutans, Downei, Other, or Unknown. We then calculated the proportion of reads assigned to species falling in each clade out of all *Streptococcus* read counts (Figure 1A, Supplemental Table S3).

**Figure 1.**
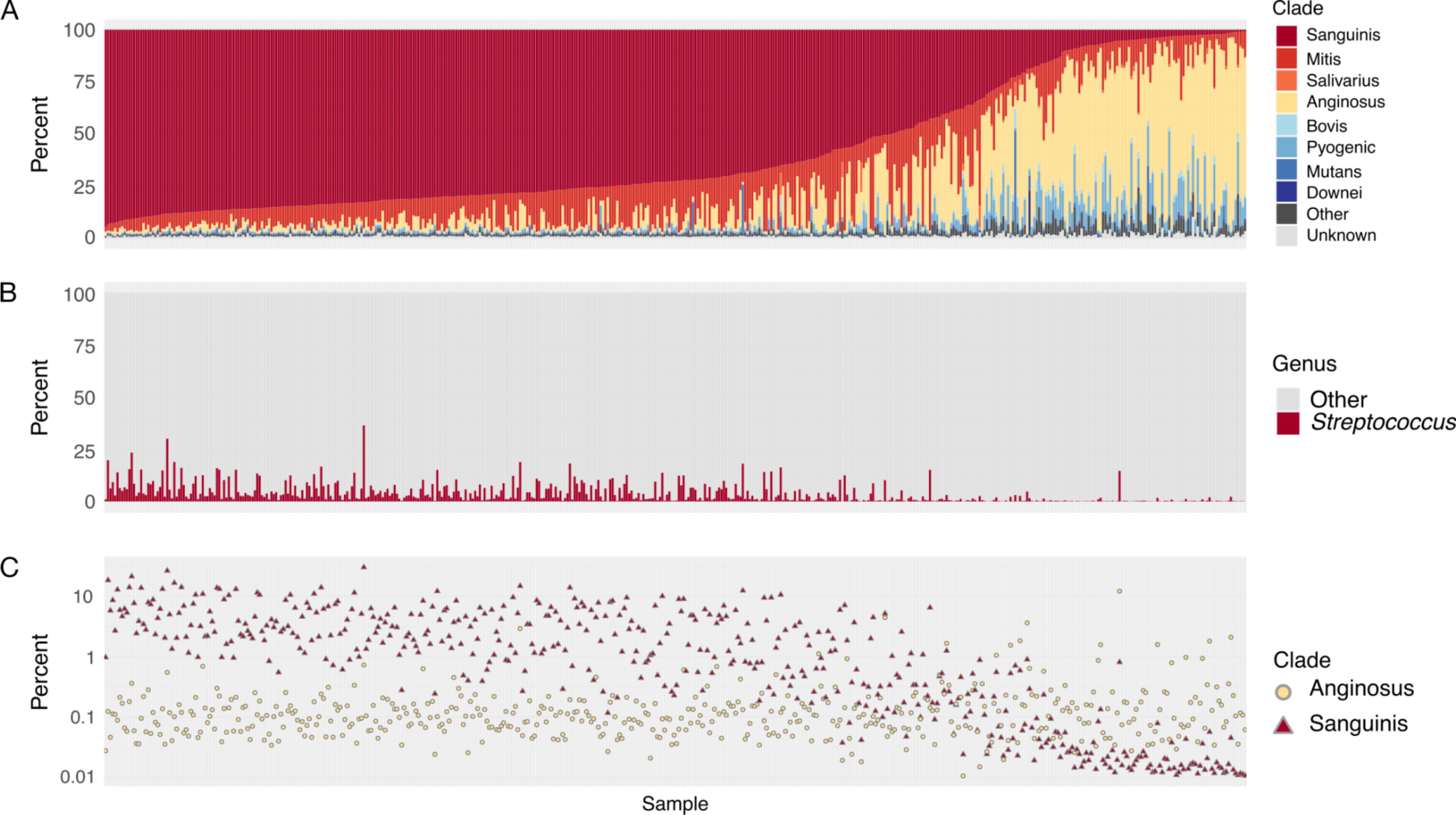
Distribution of *Streptococcus* clades in ancient dental calculus samples. **A.** Percent of *Streptococcus* reads that were assigned to each clade out of all reads assigned to *Streptococcus*, ordered by decreasing abundance of Sanguinis clade and increasing abundance of Anginosus clade. **B.** Percent of reads assigned to species in the genus *Streptococcus* and to all other genera. **C.** Percent of reads assigned to species in the Sanguinis and Anginosus clades out of all species-level read assignments.

We found that dental calculus in our global, deep-time dataset, regardless of age (Supplemental Figure S6), replicates the same pattern of *Streptococcus* clade abundance first described in^6^, with the majority having most species falling in the Sanguinis clade, while a small number of samples (71/483, 14.7%) have primarily species falling in the Anginosus clade. A Spearman’s correlation test confirmed a significant negative correlation between the two clades (**ρ** = −0.76, p < 0.0001) in this dataset. This pattern was further replicated in each dataset individually, demonstrating that this *Streptococcus* species profile is a characteristic of human ancient dental calculus generally and not the result of biases in laboratory processing, and is moreover not restricted to a particular geographic region or time period. An exception, however, was observed in a dental calculus dataset from Oceania^8^ (Supplemental Figure S7), in which the majority of samples have streptococcal species that fell within the Anginosus clade. The microbial community profile of these samples was shown to fall within the known global species variation of ancient calculus, but their microbial diversity was also distinct, likely due to the presence of as-yet unidentified taxa.

We next tested whether there was a difference in the relative abundance of *Streptococcus* overall in samples at either end of the clade spectrum (i.e., between samples with highest Sanguinis proportions and samples with highest Anginosus proportions). We found that samples with high proportions of Sanguinis species typically had a high relative abundance of *Streptococcus* overall in the dental calculus, while samples containing predominantly Anginosus clade species had a very low relative abundance of *Streptococcus,* similar to the chimpanzee calculus samples in Fellows Yates, et al.^6^ (Figure 1B). We then tested whether the samples with high proportions of Anginosus clade species show this pattern due to a loss of Sanguinis clade species, or due to an increase in abundance of Anginosus clade species, and found that it is due to a loss of Sanguinis clade species (Figure 1C), which also explains the overall lower proportion of *Streptococcus* in these samples.

We additionally investigated the abundance of several other genera with associations to *Streptococcus* described in the literature^9^. As other species in addition to *Streptococcus* may act as early and intermediate dental plaque colonizers (e.g., *Actinomyces*, *Neisseria*, *Veillonella*, *Corynebacterium*, *Capnocytophaga*, *Fusobacterium*)^10,11^, we investigated whether these taxa have higher relative abundance in samples with low *Streptococcus* abundance. None of the other early colonizing genera we investigated showed this pattern however (Supplemental Figure S8), and most had abundance patterns with a moderately strong positive correlation to that of *Streptococcus* (Spearman **ρ** = 0.43-0.80, p < 0.0001), suggesting that the physiologic conditions of these calculus biofilms did not support growth of the predominantly aerotolerant early colonizer taxa. *Actinomyces* was the only genus that was uniformly abundant across all ancient dental calculus samples, suggesting that it may play a foundational role in biofilm formation that is minimally impacted by actions of *Streptococcus*. Further, the abundance of two late colonizer genera associated with *Streptococcus, Porphyromonas*^9,12–14^ and *Methanobrevibacter*^15^, showed only weak correlations with the abundance of *Streptococcus* (Spearman **ρ** = 0.12, −0.17, respectively, p < 0.0001)*. Methanobrevibacter* abundance in particular varied widely across all samples (Supplemental Figure S9).

### Streptococcus *clade abundance is not associated with dental health, time period, geography, or processing laboratory*

We next addressed whether the difference in relative abundance of *Streptococcus* and proportions of Sanguinis and Anginosus clade species is correlated with dental pathology, laboratory processing, or sequencing outcomes. In living populations, dental health is the strongest factor associated with altered oral microbiome profiles^16^, although this does not seem to be true of ancient dental calculus microbiome profiles^17,18^. We used a large metadata-rich dataset from a single cemetery in Middenbeemster, the Netherlands^17^, which was used for a defined period of time, restricting variation due to geography and sample age.

The dental calculus in this dataset showed the same pattern of *Streptococcus* clade abundances that we observed in our global dataset (Figure 2A-C), and we found no strong correlations (correlation >= 0.4, p < 0.01) between the abundance of Sanguinis or Anginosus clade species with any of the laboratory extraction metrics, sequencing outcomes, dental records, or pathology (Figure 2D). Instead, we found that the abundance of Sanguinis clade species was strongly correlated with the PC1 and PC2 loadings in a PCA, as well as with mean library GC content, and abundance of Anginosus clade species (Figure 2D). While the correlation between the relative abundance of Sanguinis and Anginosus clade species is negative, the correlation between Sanguinis clade species abundance and mean GC content is weakly positive (Supplemental Figure S12).

**Figure 2.**
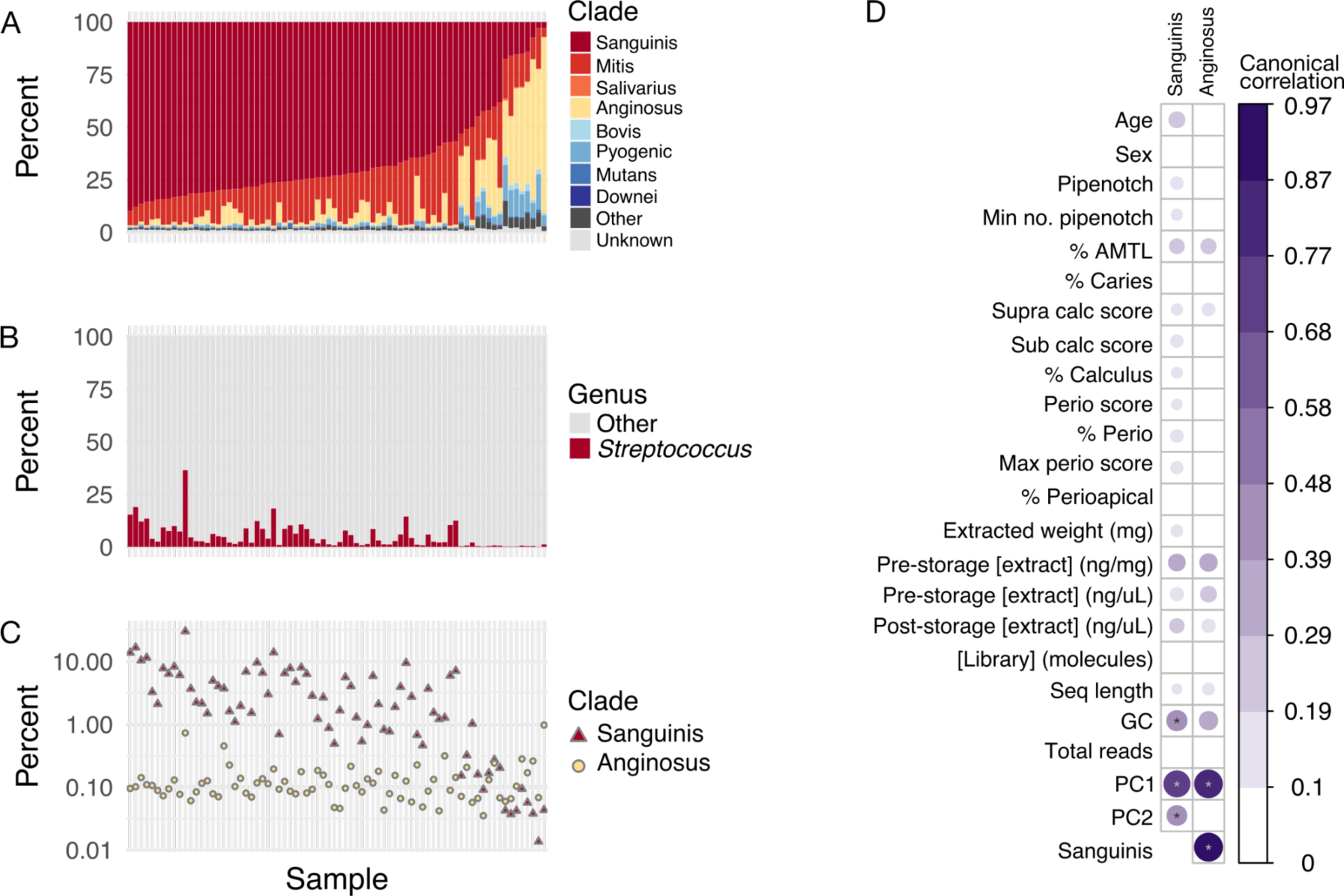
Distribution of *Streptococcus* clades in ancient dental calculus samples from Middenbeemster, the Netherlands and canonical correlations with sample parameters. **A.** Percent of *Streptococcus* reads that were assigned to each clade out of all reads assigned to *Streptococcus*, ordered by decreasing abundance of Sanguinis clade and increasing abundance of Anginosus clade. **B.** Percent of reads assigned to species in the genus *Streptococcus* and to all other genera. **C.** Percent of reads assigned to species in the Sanguinis and Anginosus clades out of all species-level read assignments. **D.** Canonical correlations between the percent of Sanguinis clade or Anginosus clade (from **A**) and archaeological metadata, laboratory, and sequencing metrics.

### Individual factors influence the dominant Streptococcus clade in ancient dental calculus

In addition to dental health, there are numerous other factors that could potentially affect the oral microbiome species profile. Many of these, such as diet, oral hygiene, drug use, and genetics, are currently difficult or impossible to investigate in ancient dental calculus. However, we may be able to determine if there are factors specific to an individual that influence an individual’s microbiome profile without knowing what those factors are. To this end, we assessed whether *Streptococcus* distribution patterns may be intrinsic to an individual by profiling the *Streptococcus* clades in multiple calculus samples collected from teeth across the dentition of four individuals. This dataset includes calculus samples representing nearly half of the dentition of each individual^19^. While three individuals had high relative proportions of *Streptococcus* (4.7%±4.2%) and exhibited mostly Sanguinis-dominated *Streptococcus* clade profiles, one individual showed the alternative pattern. For this individual, the overall relative abundance of *Streptococcus* was low (1%±1.8%), and approximately half of the calculus samples had low proportions of Sanguinis clade species (<50%) and relatively high proportions of Anginosus clade species (> 5%, Figure 3, Supplemental Figure S13). These results suggest that the low proportion of *Streptococcus* due to loss of Sanguinis clade species may be an individual-specific phenomenon, with a high probability of being observed in a single piece of calculus randomly selected for analysis.

**Figure 3.**
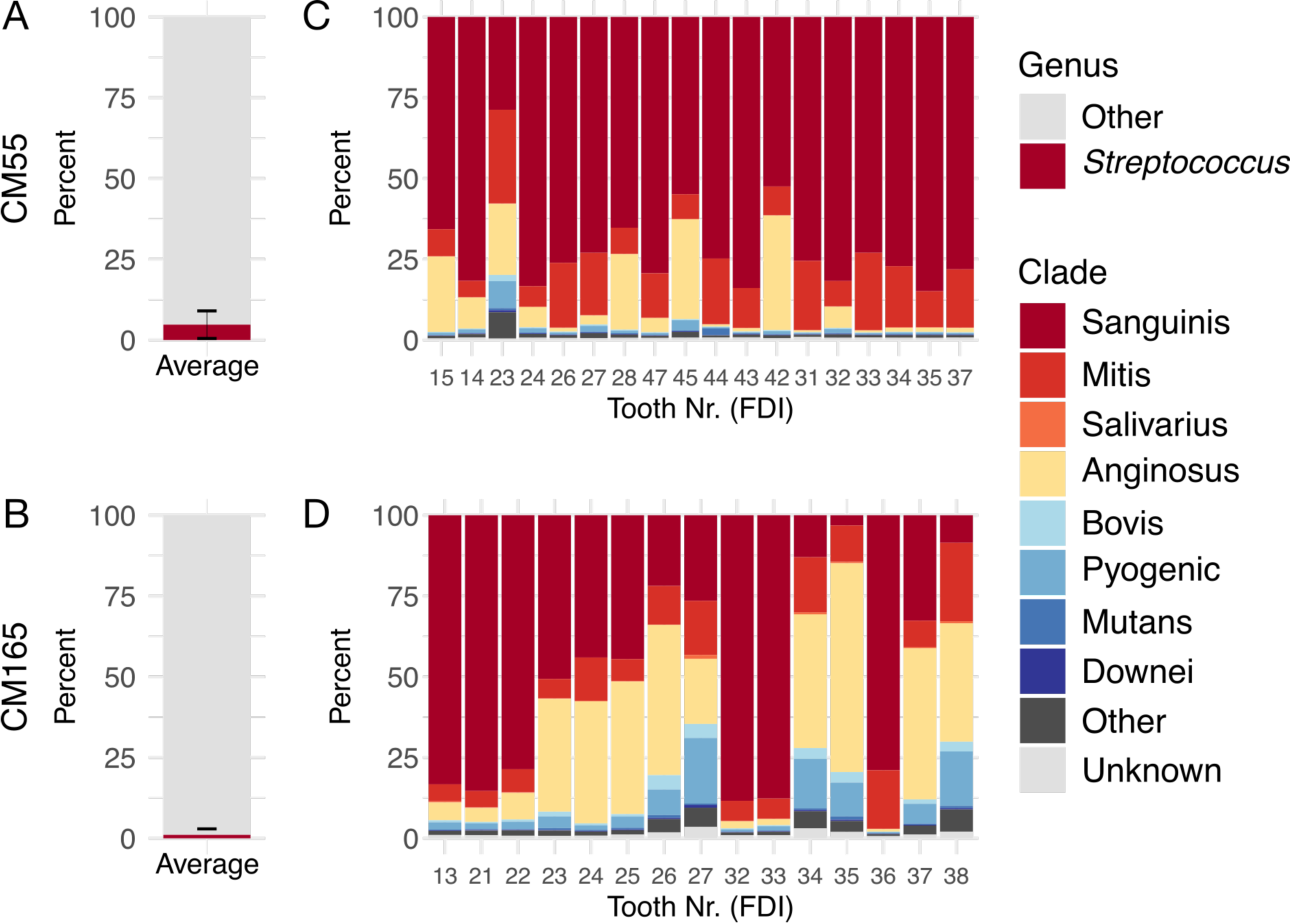
Distribution of *Streptococcus* groups in calculus of each tooth sampled from two individuals from the Chalcolithic site (ca. 4500-5000 BP) Camino del Molino, Spain. **A, B.** Average proportion of reads assigned to the genus *Streptococcus* compared to all other genera in individual CM55 (**A**) and CM165 (**B**), averaged across all teeth sampled. **C, D.** Proportion of reads assigned to species within each *Streptococcus* clade out of all reads assigned to *Streptococcus*, by tooth, in individual CM55 (**C**) and CM165 (**D**). Tooth numbers are in FDI World Dental Federation notation.

### *Global market integration/urbanization minimally impacts modern-day oral microbiome* Streptococcus *clade profiles*

While dental calculus is the only oral sample type to readily preserve in the archaeological record, there are at least seven distinct surfaces in the mouth, each of which harbors a distinct microbial community^20,21^. Oral *Streptococcus* species are known site-trophists, with different species preferentially prevalent and abundant at selected oral sites such as tongue, saliva, or dental plaque^4^. Further, because recent studies on the impacts of urbanization and industrialization have demonstrated differences in microbiomes of people living in highly urbanized, industrialized conditions relative to those living in less urbanized and industrialized locations, we next assessed whether there are differences between *Streptococcus* clade distributions in oral samples of present-day individuals living across a spectrum of urbanization and global market integration. We chose three oral sites to focus on: tooth surface (dental plaque/dental calculus), buccal mucosa (cheek swabs), and saliva, as there are publicly available microbiome datasets for these sites from groups with differing levels of urbanization/global market integration.

The tooth surface samples, calculus and plaque, contained predominantly species from the Sanguinis and Mitis clades, and they had notably higher proportions of Mitis clade species than ancient dental calculus (Figure 4A, Supplemental Table S3). Calculus samples have slightly higher proportions of Sanguinis than Mitis clade species compared to plaque; however, none of the plaque or calculus samples lacked Sanguinis clade species, in contrast to what we observed in ancient dental calculus (Figure 4C). A small number of plaque samples from individuals with low urbanization/industrialization lifestyles have high proportions of *S. mutans*, as previously noted^22^, which is due to a rise in relative abundance of this species and not a drop-out of other clade species.

**Figure 4.**
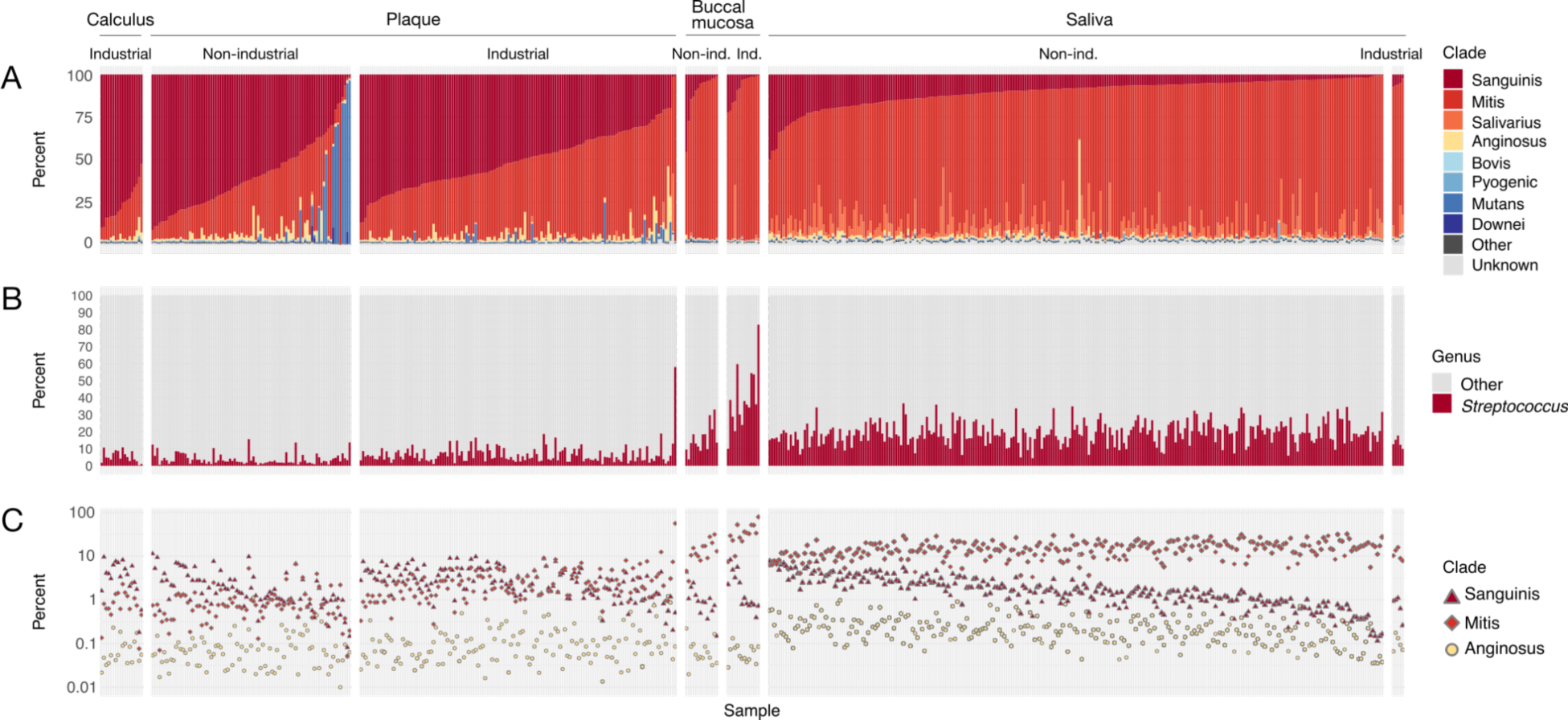
Distribution of *Streptococcus* clades in modern oral samples of calculus, dental plaque, buccal mucosa, and saliva. **A.** Percent of *Streptococcus* reads that were assigned to each clade out of all reads assigned to *Streptococcus*, ordered by decreasing abundance of Sanguinis clade and increasing abundance of Mitis clade. **B.** Percent of reads assigned to species in the genus *Streptococcus* and to all other genera. **C.** Percent of reads assigned to species in the Sanguinis, Mitis, and Anginosus clades out of all species-level read assignments.

The buccal mucosa and saliva samples have higher relative abundances of *Streptococcus* than dental plaque or calculus (Figure 4B), and they contain predominantly Mitis clade species. There were no substantial differences in the clade abundances between low and high levels of urbanization/market integration in plaque or saliva samples, although there is overall slightly lower relative abundance of *Streptococcus* in samples from low urbanization/market integration samples (Figure 4B). Buccal mucosa samples from highly industrialized contexts have much higher relative abundance of *Streptococcus* than from low industrialization contexts, yet the clade proportions remain consistent. The relative abundance of other early colonizer taxa is distinct by oral site (Supplemental Figure S10) and not clearly related to the relative abundance of *Streptococcus*. *Capnocytophaga* was the only genus with a significant difference in abundance and large effect size (p < 0.001, effect size = 0.55) between industrial plaque (10% ± 5.2%) and non-industrial plaque (3.7% ± 2.9%), suggesting it may play a more important role in structuring dental biofilms early in development rather than later.

### Species-level Streptococcus distributions show minor differences by global market economy integration

To investigate which species were dominant at each oral site within the Sanguinis, Mitis and Anginosus clades, and whether dominant species differed between samples from high and low industrialized/urbanized contexts, we created heatmaps with the relative abundance of all species in these three clades that were detected in any of the samples (Supplemental Figure S14, S15). In ancient dental calculus and modern non-industrial plaque samples where Sanguinis was the dominant clade, *S. sinensis* was the most abundant Sanguinis clade species (282/361 - 78%, Figure 5A, and 32/67 - 48%, Figure 5B, respectively), which was unexpected given that we had previously noted that *S. sanguinis* is the most abundant Sanguinis clade species in these samples. In the ancient dental calculus samples in which the Anginosus clade was more abundant, *S. constellatus* was the most abundant species in this clade (97/109, 89%).

**Figure 5.**
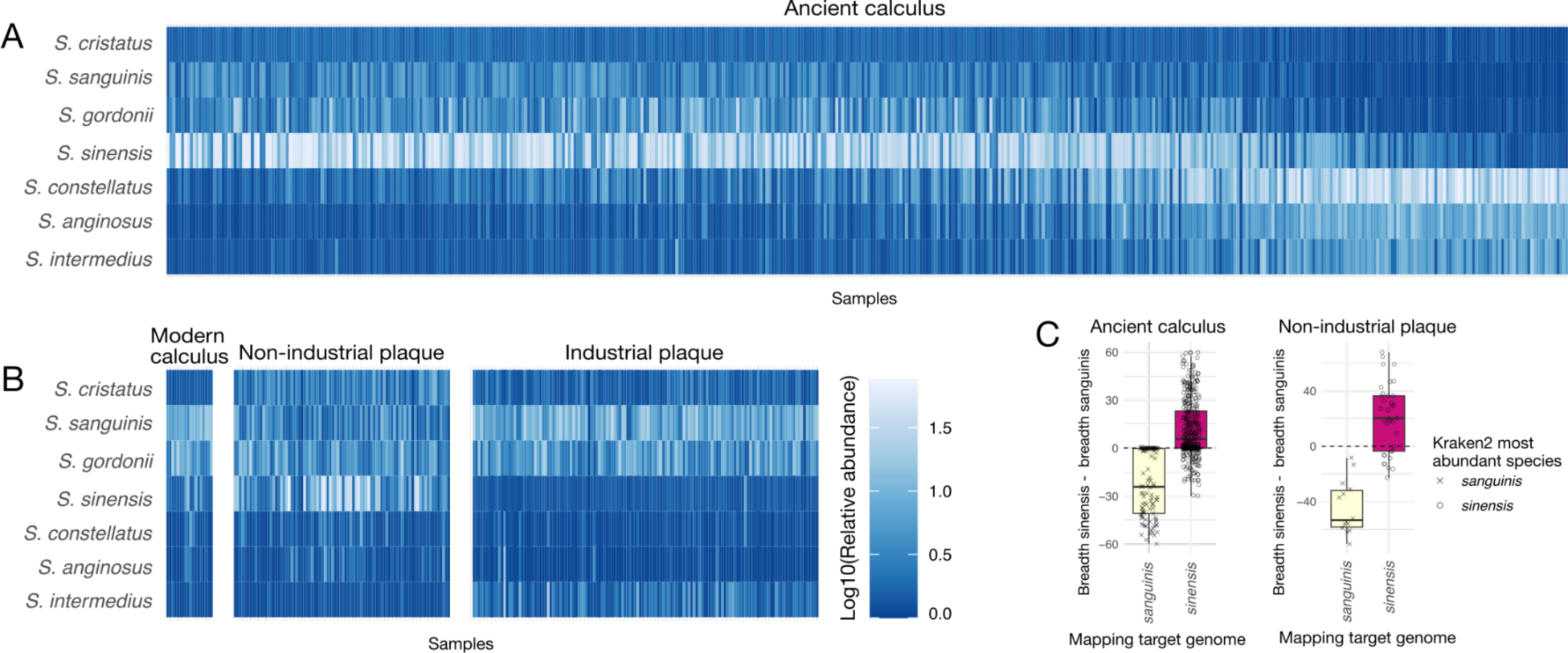
Relative abundance of Sanguinis and Anginosus clade species in tooth-adherent oral microbiome samples. Color scale is log10 of the percent relative abundance. **A.** Ancient dental calculus. Sample order is identical to Figure 1. **B.** Modern dental calculus and modern dental plaque. Sample order is identical to Figure 4. **C.** Difference in the breadth of coverage (minimum 1X depth) of *S. sanguinis* and *S. sinensis* genomes in ancient dental calculus and modern non-industrial dental plaque. Shapes indicate the *Streptococcus* species that was most abundant in each sample based on profiling with Kraken2 using the GTDB r202 database.

In modern oral samples, the most abundant species were largely consistent between high and low industrialization/urbanization samples for each oral site, with notable exceptions in plaque. Many non-industrial plaque samples have higher proportions of the Sanguinis clade species *S. sinensis* and *S. cristatus* than do any of the industrial plaque samples. Conversely, the industrialized plaque samples have higher relative abundance of the Sanguinis clade species *S. sanguinis* and the Anginosus clade species *S. intermedius* than many non-industrial plaque samples. In contrast, *S. oralis_S* is more abundant in industrial plaque than non-industrial plaque (Supplemental Figure S15). Because of the notable difference in relative abundance of *S. sinensis* and *S. sanguinis* between different human sample types, these two species appear to fulfill distinct roles in biofilm establishment and growth, such that their abundance is linked to the biofilm developmental stage, with *S. sanguinis* abundant in early-stage biofilms, but being overtaken by *S. sinensis* as the biofilm grows and matures.

### Streptococcus sinensis abundance and distribution in dental plaque and calculus

*Streptococcus sinensis* has been infrequently reported in dental plaque studies to date, and was missed in our earlier ancient dental calculus taxonomic profiling^6,8,17,18^ because it was not included in the database we used. We used two steps to confirm the presence of *S. sinensis* in the samples. First, we determined which *Streptococcus* species was the most abundant in the Kraken2 taxonomic profile for each sample, then we used genome mapping to confirm the assignment of *S. sinensis* (Figure 5C, Supplemental figure S16, S17). After mapping all dental calculus datasets against genomes for *S. sanguinis* and *S. sinensis*, we found that Kraken2 read assignments correlated well with mapping breadth and depth of coverage for each species in a majority of samples; however, for a small number of samples we observed greater mapping to the alternative reference genome, suggesting that the most abundant Sanguinis clade species in these samples is a closely-related, not-yet-described species. Future assembly of MAGs from these samples may help resolve the identity of the highly abundant Sanguinis clade species found in these samples; however, at present, MAG assembly and binning remains challenging for *Streptococcus*^23^.

### Sanguinis clade species are minimally represented in non-human primate oral microbiomes

We next assessed whether non-human primates have distinctive distributions of *Streptococcus* clades compared to humans by examining the species present in calculus and oral swabs. The overall abundance of *Streptococcus* varied substantially across non-human primates and oral sites, with *Streptococcus* species being generally lower in dental calculus than in oral swabs (Figure 6). Chimpanzees had among the lowest relative abundance of *Streptococcus* in both dental calculus (0.33% ± 0.23%) and oral swabs (3.1% ± 1.6%), while *Streptococcus* made up more than half of the species identified in oral swabs from vervet monkeys (59% ± 20%). Each non-human primate species had a distinct distribution of *Streptococcus* clades, and many of the most abundant *Streptococcus* species did not fall into previously described clades (Figure 6A, Supplemental Table S3), particularly in the oral swab samples. A PCA of beta-diversity differences highlights how the abundance of particular clades of *Streptococcus* may contribute to overall diversity differences between hosts and oral sites (Figure 7, Supplemental Figure S18).

**Figure 6.**
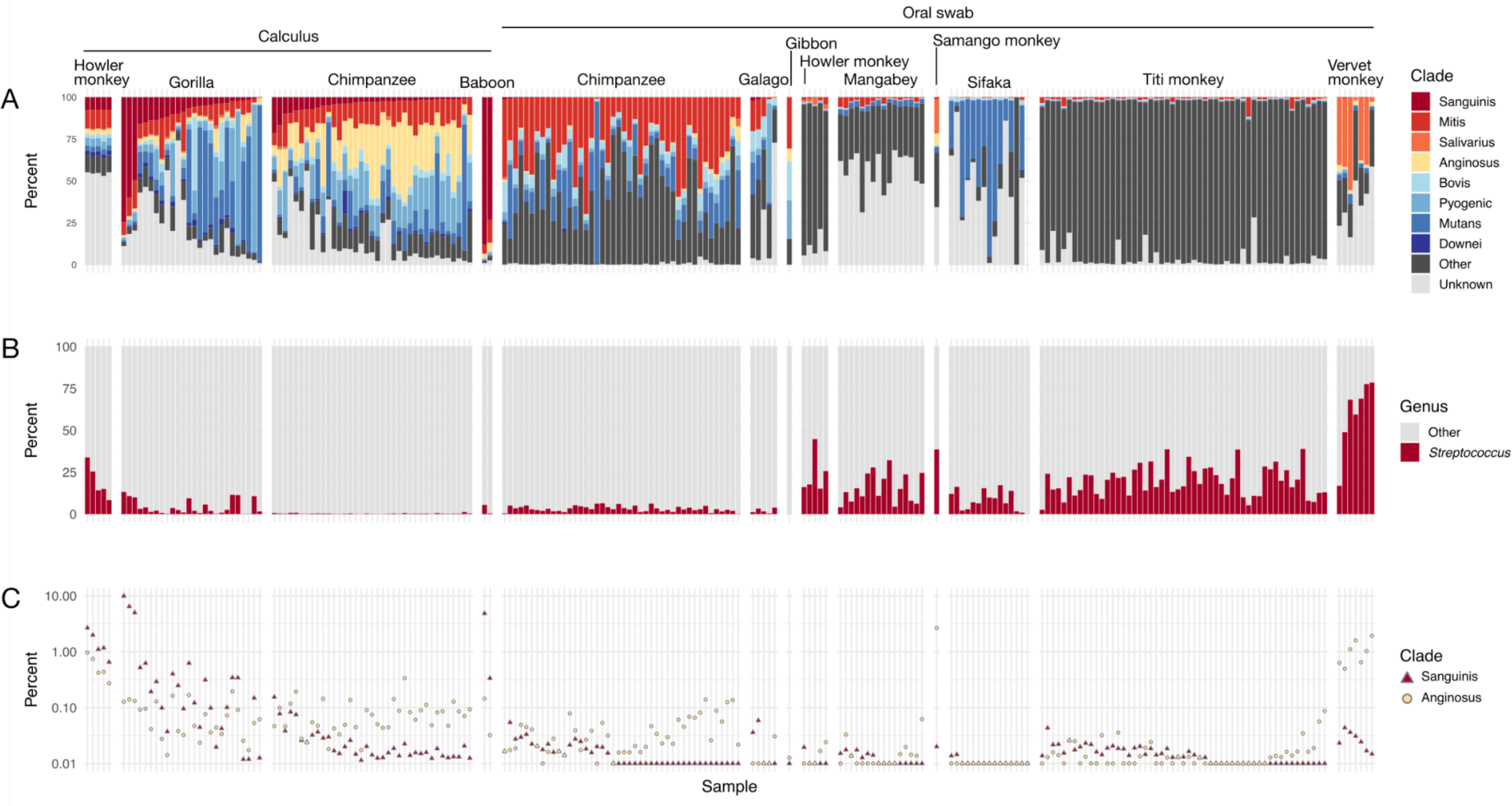
Distribution of *Streptococcus* clades in dental calculus and modern oral swabs of non-human primates **A.** Percent of *Streptococcus* reads that were assigned to each clade, ordered by decreasing abundance of Sanguinis clade and increasing abundance of Mitis clade. **B.** Percent of reads assigned to species in the genus *Streptococcus* and to all other genera. **C.** Percent of reads assigned to species in the Sanguinis, Mitis, and Anginosus clades out of all species-level read assignments.

**Figure 7.**
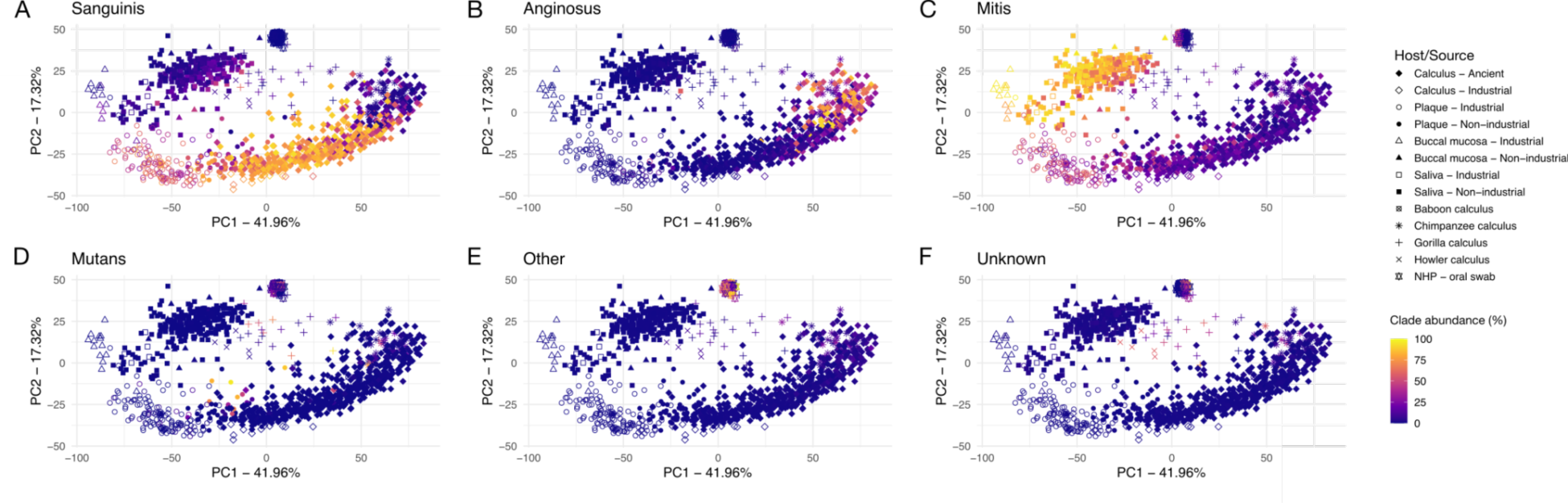
Principal components analysis plot of ancient and modern human and non-human primate oral microbiomes. Shapes indicate sample host, colored by proportion of each clade out of all *Streptococcus* clades: **A.** Sanguinis clade, **B.** Anginosus clade, **C.** Mitis clade, **D.** Mutans clade, **E.** Other clades, **F.** Unknown clades.

In contrast to humans, only 5 of 107 non-human primate dental calculus samples (∼5%) had a predominantly Sanguinis clade profile: three gorilla samples and two baboon samples. While this represents a minority profile for gorillas (3 of 26), further study of baboon dental calculus is needed as only two dental calculus samples were sufficiently preserved for analysis. From these results, it appears that dental calculus dominated by Sanguinis and Mitis clades may be a particularly human feature, although further investigation of baboon samples will be necessary to confirm this. Other early colonizer genera were generally infrequently detected and low in abundance across non-human primate oral samples, with a few exceptions (Supplemental Figure S11).

### Streptococcus gene content differs between hosts and oral sites

Given the ecological diversity and distinct niches occupied by *Streptococcus* species in human and non-human oral microbiota, we tested whether there are particular genes in *Streptococcus* that are enriched or depleted in the primate hosts and oral sites. This includes genes that are significantly enriched in human shedding surfaces (buccal mucosa/saliva) compared to non-shedding surfaces (dental plaque and calculus), between human and non-human primates for either shedding and non-shedding surfaces, and between human ancient and modern dental calculus (Supplemental Table S5). A total of 2,615,774 genes attributed to *Streptococcus* were identified across all samples. The majority of genes that we found significantly enriched between groups were genes of unknown function (Supplemental Table S5), which limits the conclusions we can draw regarding functional specificities of *Streptococcus* in different hosts and at different oral sites. This highlights the necessity of continued laboratory functional characterization of host-associated microbes to improve our understanding of microbial metabolic functioning and the impacts it has on both the microbiome community and the host.

## Discussion

The highly diverse genus *Streptococcus* shows distinct host and site-tropism within the oral cavities of primates. While these differences may reflect host differences in salivary composition^24^ and dietary differences among primate species^6^, human-associated *Streptococcus* appear to a small extent to also be affected by human hygiene practices and level of global market economy integration/urbanization. Humans appear to be uniquely enriched in species from the Sanguinis and Mitis clades, which are otherwise rare in non-human primates with the possible exception of baboons. As the savannah territory and tuber consumption of baboons is hypothesized to be similar to that of early humans, the ability of Sanguinis and Mitis clade species to utilize dietary starch may have provided an ecological advantage that lead to the dominance of these species in the mouths of starch-consuming primates^6^.

Using a large and diverse dataset of more than 400 well-preserved dental calculus samples, we were able to confirm that human ancient dental calculus can be grouped into two categories based on *Streptococcus* abundance profiles, which are consistently found across time and geography. By using publicly available data that was produced in different labs using different DNA extraction and library build techniques, we also demonstrate that this observation is robust to laboratory processing methods, and is likely a true biological phenomenon. The factors driving these distinct ecological profiles are not yet known, and although we found no associations between streptococcal profiles and osteological measures of oral pathology, we cannot rule out that there may be associations with specific immune responses or soft tissue pathology, which cannot be measured by DNA analysis or osteological examination of archaeological remains.

However, other host physiology-related explanations seem likely drivers of different *Streptococcus* clade profiles. For example, early studies of *in situ* biofilm formation on tooth surfaces reported two distinct groups of study participants that developed plaque at different rates^25–27^. Further work found differences in salivary properties and composition between “rapid” and “slow” plaque-forming participants that influenced *Streptococcus* species^28–30^. We speculate that the *Streptococcus* profiles of ancient dental calculus may reflect the “rapid” and “slow” plaque formation documented in these early plaque development studies; however, whether there is a relationship between the two would require further *in situ* or *in vitro* modeling to understand.

The near absence of early colonizer species other than *Actinomyces* in dental calculus with low overall *Streptococcus* abundance suggests that those biofilms may have a distinct pattern of species acquisition and turnover that differs from the well-described oral biofilm succession model^31^. In ancient dental calculus, we found moderate to strong correlations between the abundance of *Streptococcus* and of most early colonizer species we examined, which was particularly striking for *Corynebacterium* and *Capnocytophaga*, which are important species for structuring the dental plaque biofilm^9,32^ and are abundant across non-industrial and industrial dental plaque samples.

*Capnocytophaga* was on average less abundant in non-industrial plaque samples than industrial plaque samples, which hints that overall plaque biofilm species and structural turnover to a more anaerobic, mature community may be related to a loss of this genus, given that the taxonomic profile of these samples appears intermediate between ancient dental calculus (which represents fully mature dental biofilms) and dental plaque from industrialized populations (which represents earlier stage dental biofilm formation due to toothbrushing). The low abundance of *Capnocytophaga* in gorilla and chimpanzee calculus suggests that it may be particularly important in structuring human plaque biofilms, despite being a core member of the primate oral microbiome^6^. As *Actinomyces* are uniformly abundant in the tooth-adherent biofilm samples we examined, including ancient and modern human dental calculus, human dental plaque, and non-human primate dental calculus, this genus may be the ancestral early colonizer for dental biofilms, with humans later adding *Streptococcus*, with concomitant changes in the taxa that successively colonize the biofilm and now differentiate human and non-human primate dental biofilms.

In previous work, we reported *S. sanguinis* to be the most abundant *Streptococcus* species in human dental calculus^6,17,18,22^, but here we find that while *S. sanguinis* is present, *S. sinensis* is actually the most abundant Sanguinis clade species present in both ancient dental calculus and modern non-industrial plaque samples. This discrepancy is due to the fact that the custom RefSeq database used for taxonomic classification in prior studies did not include any *S. sinensis* genomes, and reads from this species were likely mis-assigned to the closely-related *S. sanguinis* instead, a known phenomenon in taxonomic assignments using incomplete databases^33^. Inconsistencies in NCBI taxonomy may also account for the absence of *S. sinensis* in taxonomic profiles, such as with the NCBI genome *Streptococcus* sp. DD04, which has been reclassified as *S. sinensis* by GTDB. Despite their close phylogenetic relationship, *S. sinensis* and *S. sanguinis* appear to thrive in different biofilm environments.

*Streptococcus sinensis* was originally isolated from a patient with infective endocarditis^34^ and was later found to be part of the oral microbiome^35^, but its physiology and biochemistry have not yet been extensively explored^36^. Although it was not included in the phylogeny presented in Richards, et al. (2014)^3^, our clustering of the type strain genome by average nucleotide identity placed it in the Sanguinis clade, with high similarity to *S. cristatus*, reflecting the phylogenetic placement of the species reported by others^37,38^. The high prevalence and abundance of *S. sinensis* in both ancient dental calculus and modern plaque from populations with low global market integration, but not in plaque from highly industrialized populations suggests that *S. sinensis* may therefore represent the first described VANISH (volatile and/or associated negatively with industrialized societies of humans) taxon of the human oral microbiota^39,40^.

The high abundance of *S. sinensis* in ancient and non-industrialized oral biofilms suggests that the species prefers more mature, anaerobic biofilm environments. This pattern contrasts with the pattern observed in dental plaque from heavily industrialized populations, in which early biofilm colonizer streptococcal species preferring more aerobic environments, such as *S. sanguinis* and *S. gordonii*, predominate. Regular and/or frequent toothbrushing to remove dental plaque in heavily studied industrial populations may prevent the oral biofilm from maturing to an anaerobic, reduced state that can support growth of *S. sinensis*, potentially explaining why it is not commonly reported in dental plaque of heavily industrialized populations. Dental hygiene, in particular regular tooth brushing, has been proposed to account for differences in species prevalence and abundance between dental plaque and ancient dental calculus as well as between dental plaque from populations with high and low global market integration and urbanization^17,18,22^, and may be one of the strongest factors shaping oral microbiome composition today.

Certain genetic differences between *S. sinensis* and. *S. gordonii/sanguinis* may explain the differences in abundance between these species in ancient dental calculus. Nearly all sequenced genomes of *S. gordonii* and *S. sanguinis* contain a gene encoding the protein AbpA that allows them to bind human salivary amylase^6^. Expression of this protein could offer a colonization advantage to *S. gordonii* and *S. sanguinis*, as salivary amylase is part of the enamel pellicle that forms the base layer on which dental plaque biofilms grow^41^. This protein was suggested to play a role in shaping the human-specific dental plaque biofilm^6^, and the near ubiquity of the gene in sequenced genomes of *S. gordonii* and *S. sanguinis* suggests it plays an important role in the physiology of these species. In addition, many *S. gordonii* and *S. sanguinis* genomes contain *gspB* or a homologue (*hsa, srpA*) that may play a role in substrate binding or nutrient acquisition, as it allows binding to sialic acids^42^, which are abundant on salivary mucins. In contrast, none of the four *S. sinensis* genomes in NCBI contain *abpA* or *gspB/hsa/srpA*, suggesting *S. sinensis* does not share the colonization advantage of *S. gordonii* and *S. sanguinis* for early biofilm formation. The high relative abundance of *S. sanguinis* in dental plaque from industrial populations that practice regular tooth brushing, which represents early-stage dental plaque biofilms, and the contrasting high relative abundance of *S. sinensis* in ancient dental calculus and dental plaque of groups with low global market economy integration, which represent more mature dental plaque biofilms, supports the preference of these *Streptococcus* species for different stages of biofilm development.

An inverse relationship between the abundance of *Streptococcus* and *Methanobrevibacter* has been reported in ancient dental calculus^17^, which is argued to reflect the overall oxygen tolerance of other abundant species in these samples. Although we see this trend in our large ancient dental calculus dataset here, *Methanobrevibacter* abundance is highly variable between samples, ranging from 0% to nearly 40% even across samples in which *Streptococcus* is nearly absent, and the correlation between the genera is weak. *S. constellatus* has been reported to support *Methanobrevibacter* growth^15^, and this is the Anginosus clade species that is most abundant in the dental calculus with low *Streptococcus* abundance, perhaps providing support for a metabolic interdependence between the two. However, the ecological role of *Methanobrevibacter* in shaping oral communities with low *Streptococcus* abundance may also be filled by other taxa in its absence, calling into question the importance of *Methanobrevibacter* itself in shaping biofilm community structure^43^, and emphasizing instead a set of as yet undefined metabolic features that may be shared by numerous taxa.

Despite high genetic heterogeneity and horizontal gene transfer within the oral streptococci, species-specific preferences for distinct oral niches appear to be relatively consistent across time, geography, and cultural practices. The role oral streptococci fill in dental plaque biofilm development appears to strongly affect the mature biofilm species profile. Specialization of the Sanguinis clade species within the human oral cavity concomitant with the increase in starch in human diets may have been critical step in shaping the human oral microbiome profile that exists today, while the recent adoption of regular tooth brushing may be driving population-wide loss of species like *S. sinensis* from the oral microbiome. Further investigation of ancient dental calculus and oral biofilm samples from under-studied living populations is necessary to provide a better understanding of the evolutionary history of oral streptococci and how changing cultural practices are impacting oral microbiome communities today. Employing an anthropological approach informed by human evolutionary biology and paleogenomics enables us to broaden our understanding of what makes a healthy, stable oral biofilm community.

## Methods

### Data selection and download

Ancient dental calculus metagenomic data from studies published prior to June 2022 including more than 2 samples that were Illumina shotgun sequenced, and were not explicitly used for extraction or decontamination method testing were downloaded from the European Nucleotide Archive (ENA)^6–8,17,18,44–51^. All samples are listed on the Ancient Metagenome Directory^52^. Modern human^6,18,21,22,53–55^ and non-human primate^6,56–60^ Illumina shotgun sequenced data were likewise downloaded from the ENA. A list of all samples and accessions is in Supplemental Table S1. This resulted in a starting dataset of 541 ancient human calculus samples, 537 modern human samples (18 calculus, 220 plaque, 28 buccal mucosa, 271 saliva), 107 dental calculus samples from non-human primates (chimpanzee, gorilla, baboon, howler monkey), and 197 oral swabs from modern non-human primate (chimpanzee, galago, gibbon, howler monkey, mangabey, samango monkey, sifaka, titi monkey, vervet monkey).

### Data processing

Raw fastq files for all samples were processed with the nf-core/eager pipeline^61^. Settings were left in default except for bwa on ancient samples, for which the following flags were used −l 32 -n 0.01. All samples regardless of host species were mapped against the human genome, and all reads that mapped were discarded from downstream analysis. All remaining reads were taxonomically classified using Kraken2^62^ and the Genome Taxonomy Database (GTDB)^63–65^ r202 database provided on the Struo2 ftp server^66^. Metaphlan-formatted output tables were joined using the KrakenTools^67^ script combine_mpa.py. Differences in the proportion of reads that were assigned taxonomy are not related to the sequencing depth (Supplemental Figure S1). We confirmed that the *Streptococcus* species profile was similar to that published by Fellows Yates, et al. (2021)^6^ (Supplemental Figure S5), and placed the *Streptococcus* genomes found in the GTDB r202 database but not the custom RefSeq database used by Fellows Yates, et al. into clades by clustering based on ANI with dRep^68^ (Supplemental Table S2, for details see the Supplemental Methods).

### Sample preservation assessment

Preservation of all samples was assessed with the R package cuperdec^6^. Samples were grouped as non-human primate, ancient human calculus, or modern human oral, and preservation cut-offs were determined individually for each of the three groups (Supplemental Figures S2-S4). All samples that were determined to be poorly preserved were discarded from downstream analysis (Supplemental Table S1). This left 482 ancient human samples, 532 modern human samples (18 calculus, 220 plaque, 28 buccal mucosa, 267 saliva), 70 ancient primate calculus samples, and 147 modern primate oral swabs.

### Assessment of Streptococcus clade distributions

We calculated the proportion of reads from *Streptococcus* species in each of these groups out of all *Streptococcus* species-assigned reads in each sample. Further, within each sample, we calculated the proportion of reads that were assigned to any species in the genus *Streptococcus* vs. all other genus assignments. We additionally calculated the proportion of reads assigned to all species per clade out of all species assignments. Lastly, we calculated the relative abundance of all species in each sample and selected out the *Streptococcus* species for plotting in a heat map. Percentages were log10-transformed after adding a value of +1 to all percents, to better visualize the different abundances across species and samples. The relative abundance of additional taxa was calculated in the same way. Principal components analysis was performed with the R package mixOmics^69^. For details see Supplemental Methods.

### Correlations with oral pathology

Correlations between the proportion of *Streptococcus* clades and oral pathology in the historic Middenbeemster dataset were assessed with canonical correlation analysis, as performed in Velsko, et al. 2022^17^, following^70^. Input tables contained selected metadata categories (Supplemental Table S1 from source publication), as well as the proportion of Sanguinis and Anginosus clade *Streptococcus* species out of all *Streptococcus* species detected from the Kraken2 taxonomic table generated for this study. For details see the Supplemental Methods.

### Streptococcus sanguinis and S. sinensis genome mapping

Representative genomes of 9 *S. sanguinis* species from GTDB classification and the 4 available *S. sinensis* genomes were downloaded from NCBI (Supplemental Table S4) and concatenated into a single fasta file. All non-industrial plaque samples and all human ancient calculus samples were mapped to this concatenated file with bwa aln using the flags −l 32 -n 0.01. Variants were called with bcftools mpileup, and the calls were filtered for those with a quality greater than 20 and a depth of >= 2. Samples were further individually mapped against the *S. sanguinis* and *S. sinensis* genomes with the highest breadth and depth of coverage.

### Gene content enrichment

To assess differences in the gene content between sample groups, we annotated the gene content of all samples using the Global Microbial Gene Catalog (GMGC)^71^. For gene coverage normalization across samples as reads per kilobase, we used RRAP^72^. Genes attributed to *Streptococcus* based on the GMGC-provided taxonomy list were subsetted from the normalized table and used to assess differences in presence/absence of genes between groups. For details see the Supplemental Methods.

### Data visualization

All plots were generated in R with ggplot2^73^ unless otherwise noted. Plots were assembled with the R package patchwork^74^. Statistics were calculated with the R package rstatix^75^. The viridis color package^76^ was used for continuous colors.

## Data availability

All data analyzed here was published in other studies and is publicly available. All accessions are listed in Supplemental Table S1.

## Code availability

The underlying code for this study is available at https://github.com/ivelsko/oral_streptococcus_clades.

## Acknowledgments

We thank Alexander Hübner for discussion on gene content analyses. This work was supported by the Werner Siemens Stiftung (“Paleobiotechnology” to C.W.), the Deutsche Forschungsgemeinschaft (DFG, German Research Foundation) under Germany’s Excellence Strategy (EXC 2051 Project-ID 390713860), the American School for Prehistoric Research (ASPR), the Max Planck Society, and Harvard University.

## Author contributions

C.W. and I.M.V. conceived the project. I.M.V. performed the analyses and wrote the manuscript with input from C.W..

## Supplemental Material for

### Supplemental Methods

#### Comparison of MALT RefSeq and Kraken2 GTDB r202 Streptococcus profiles

Fellows Yates, et al. (2021)^1^ used MALT with a custom RefSeq database to profile the species in their ancient calculus samples. As MALT requires a substantial amount of memory to run and takes many hours per sample, and this database is now out-dated, it was not feasible to use MALT for profiling the samples in this study. We chose to use Kraken2 and the GTDB database r202 (the most recent release at the time this study was performed) because of Kraken’s speed, and the comprehensive species representation in GTDB. To confirm that the *Streptococcus* species profiles we found with Kraken2/GTDB were similar to that seen with MALT/customRefSeq, we compared the *Streptococcus* clade profiles for two datasets for which we already had MALT/customRefSeq species tables: Fellows Yates, et al.^1^, and Velsko, et al.^2^. We found the *Streptococcus* clade profiles were highly comparable (Supplemental Figure S5A-C), although there were some notable differences with the non-human primate clade distributions. This was likely due to the differences in *Streptococcus* species in each database (Supplemental FIgure S5D).

#### Assessment of Streptococcus clade distributions

To be able to group *Streptococcus* genomes that had reads assigned by Kraken2 but for which the clade was unknown (because it is unnamed, or uploaded to NCBI after the publication of Richards, et al. (2014)^3^, we ran ‘dRep cluster’ on all *Streptococcus* genomes with hits in any sample. *Streptococcus* genomes were then assigned to a clade, defined by Richards, et al. 2014, based on dRep primary clustering phylogeny. If a genome fell outside of these clades, it was assigned “Other”, while genomes that fell outside of these clades and were basal to all known/named clades were assigned “Unknown”.

#### Assessment of species distributions of additional taxa

We additionally investigated the abundance of six selected genera for their role in early biofilm colonization and development, as well as two late colonizer species that are known to interact with *Streptococcus*. The early colonizer genera were: *Neisseria*, *Actinomyces*, *Veillonella*, *Fusobacterium*, *Capnocytophaga*, and *Corynebacterium*. The late colonizer species were *Porphyromonas gingivalis* and *Methanobrevibacter oralis*. To determine the abundance of these taxa, we used the same approach as we used to calculate the proportion of *Streptococcus* vs all other genera. Within each sample, we calculated the proportion of reads that were assigned to any species in each of the above genera vs. all other genus assignments, or, for the two late colonizer species we calculated the proportion of reads assigned to each species vs. all other species assignments. We performed a Spearman correlation test to determine whether there was a correlation between the abundance of each early colonizer genus or each late colonizer species and the abundance of *Streptococcus* in each sample. P < 0.05 was considered significant.

#### Correlations with oral pathology

The function canCorPairs from the R package variancePartition^4,5^ was used to perform canonical correlations, while the cor.mtest function in the R package corrplot^6^ was used to perform statistical tests. Correlation matrix plots were generated with the function corrplot in the same package. To focus on the strongest correlations, we considered only correlations ≥ 0.4 with a significance of p ≤ 0.01 to be significant.

#### Streptococcus *genome mapping*

Representative genomes of the 9 *Streptococcus sanguinis* species from GTDB classification and the 4 available *S. sinensis* genomes were downloaded from NCBI (Supplemental Table S3) and concatenated into a single fasta file. All non-industrial plaque samples and all human ancient calculus samples were mapped to this concatenated file with bwa aln using the flags −l 32 -n 0.01. Variants were called with bcftools mpileup with the flags -Ou, bcftools call --Ou -mv - A --ploidy 1, and the calls were filtered for those with a quality greater than 20 and a depth of >= 2. The *S. sanguinis* (GCF_003943655.1) and *S. sinensis* (GCF_000767835.1) genomes with the highest breadth and depth of coverage were selected and all samples mapped against these individual genomes with bwa aln using the same parameters as above, then variants were called with bcftools in the same way as above. The vcf files were converted to tsv using bcftools norm -m and bcftools query -f ‘%CHROM\t%POS\t%REF\t%DP\t%ALT\n’ to separate multiallelic snps into one observed snp per line per site for data analysis.

In addition, we mapped all non-industrial plaque samples and all human ancient calculus samples to a file containing *S. sanguinis* (GCF_003943655.1) and *S. sinensis* (GCF_000767835.1) genomes as well as four MAGs of Sanguinis clade *Streptococcus* that were assembled from modern and ancient dental calculus in^7^, using bwa aln and the same parameters as above. This allowed us to determine if the dominant Sanguinis clade species in our samples may be one that was not represented by genomes in NCBI RefSeq or Genbank.

#### Gene content enrichment

Collapsed reads from all samples were mapped against the GMGC database GMGC10.95nr with bowtie2 and the following flags: -D 20 -R 3 -N 1 -L 20 -i S,1,0.50 --no-unal. This allowed for mapping ancient damaged reads, but was applied to all samples, both ancient and modern. The bowtie2-mapped bam files of each sample mapped against the Global Microbial Gene Catalog (GMGC) 10.95nr were used as input for RRAP^8^ for normalization by reads per kilobase, with the parameters –skip-indexing and –skip-rr. Since reads used for mapping were collapsed read pairs, we used the collapsed read fastq file as both a read 1 and a read 2 fastq file for RRAP. Three groups of samples were run individually through RRAP: the modern human oral samples, the ancient human calculus, and the non-human primate samples. This produced three output files of gene content normalized by reads-per-kilobase, one for each sample group. Significant differences in gene presence/absence between groups was assessed with a Wilcox test and effect size using the R package rstatix^9^. Prior to testing for significant differences, tables were subsetted to include only genes present in at least 30% of all samples being compared. Genes were considered significantly enriched in one group if multiple test-corrected p-values were less than 0.05 and the effect size was at least 0.4.

## Supplemental Figures

**Figure S1.**
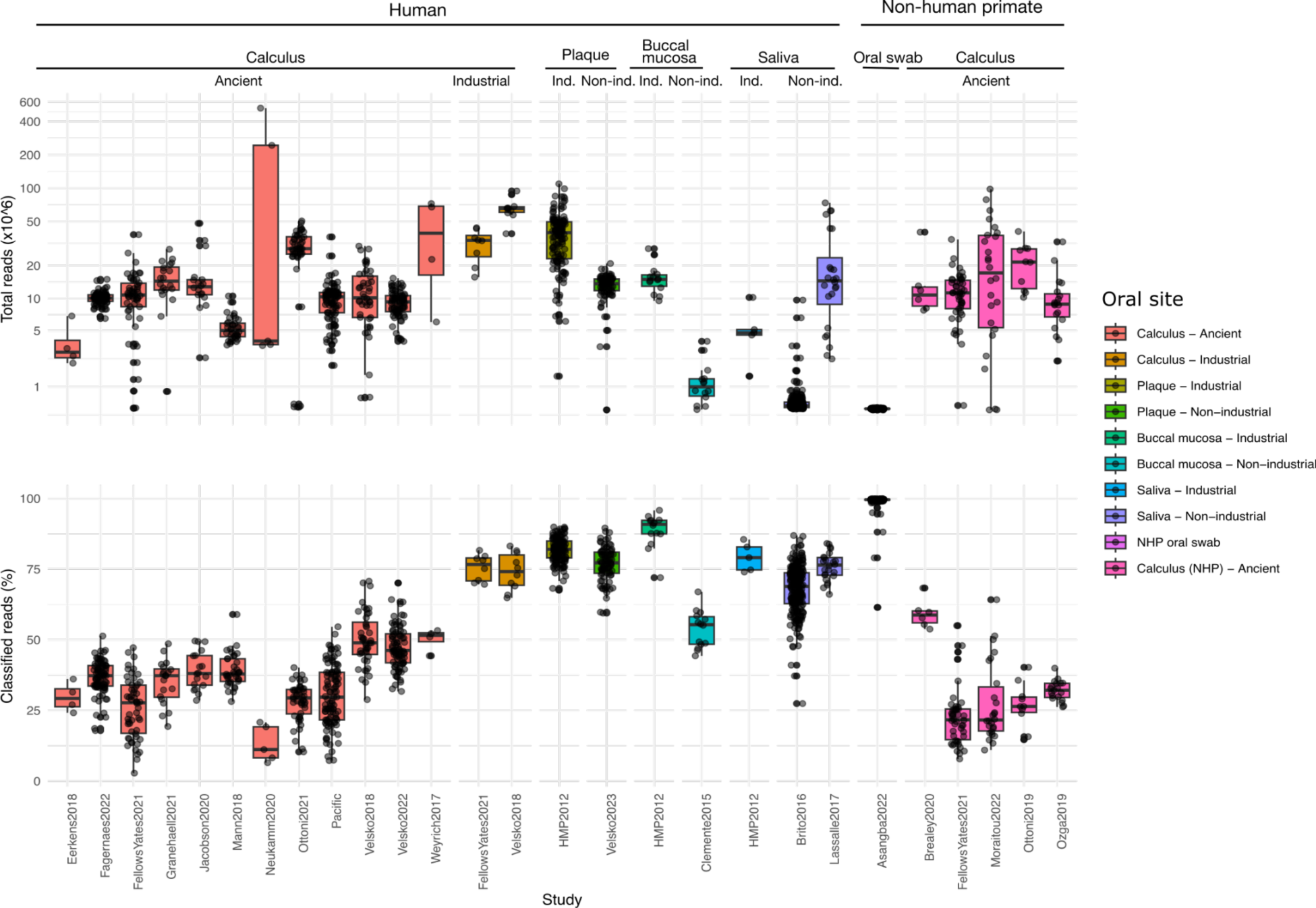
Read classification stats. **A.** Total reads in each sample grouped by study. **B.** Percent of classified reads in each sample, grouped by study. Ind. - Industrial; Non-ind. - Non-industrial.

**Figure S2.**
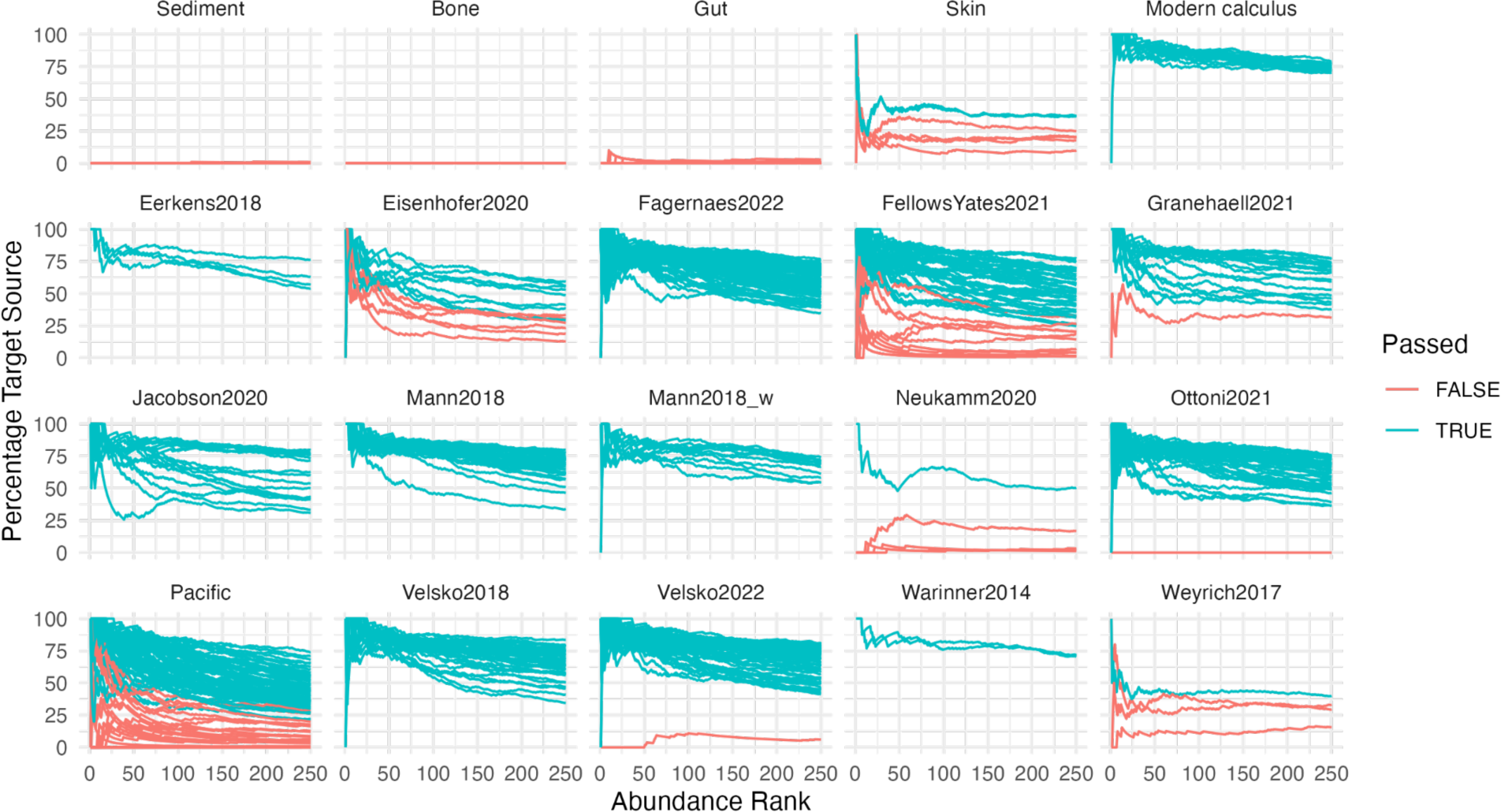
Cumulative percent decay curves for ancient dental calculus samples and environmental controls. Lines indicate the percent of all species at and above the abundance rank that are classified as oral. Passed - False indicates the sample did not pass the preservation cut-off; Passed - True indicates the sample passed the preservation cut-off and is well-preserved. All samples that did not pass were excluded from all downstream analyses.

**Figure S3.**
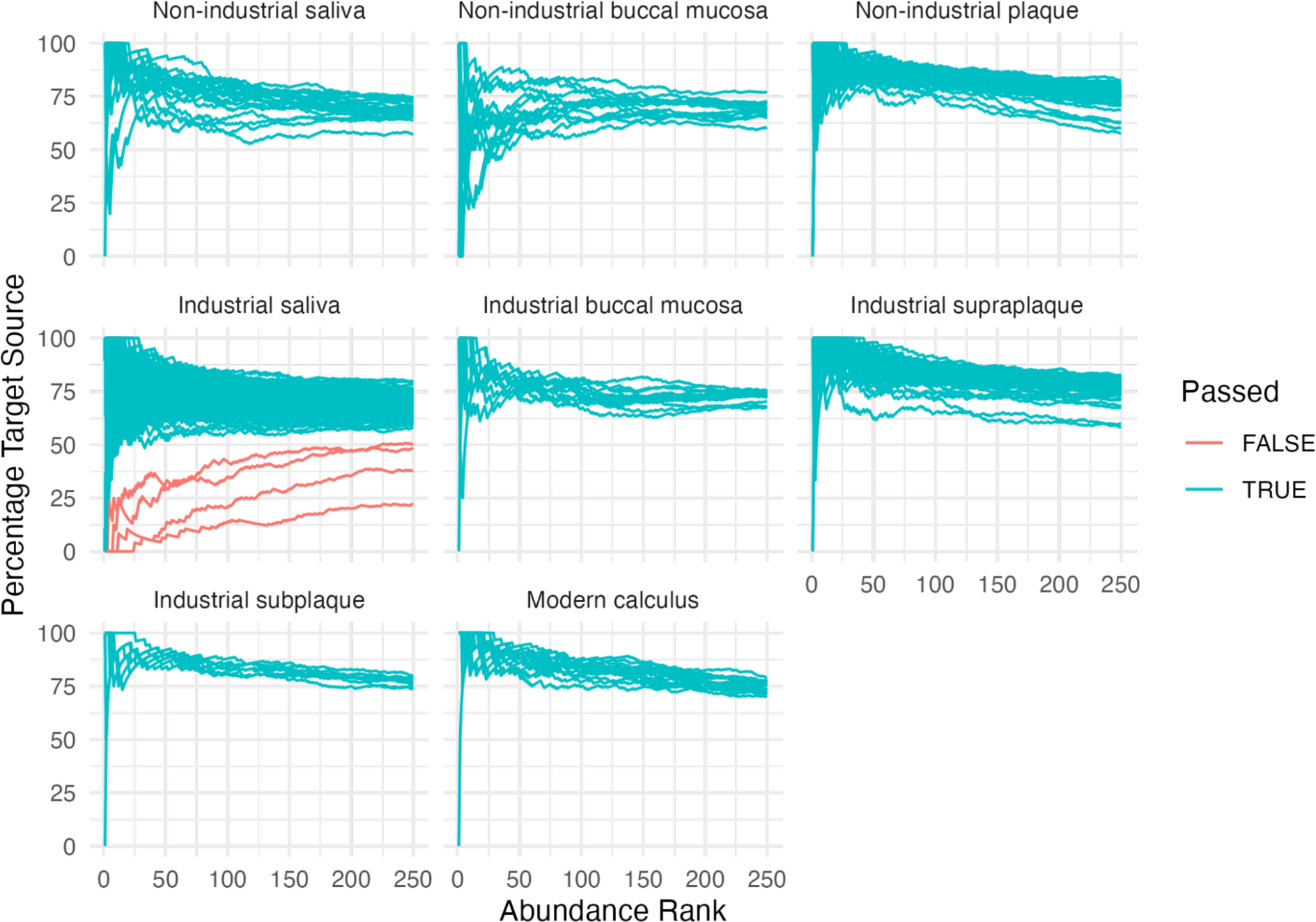
Cumulative percent decay curves for human modern oral samples. Lines indicate the percent of all species at and above the abundance rank that are classified as oral. Passed - False indicates the sample did not pass the preservation cut-off; Passed - True indicates the sample passed the preservation cut-off and is well-preserved; supraplaque - supragingival plaque; subplaque - subgingival plaque. All samples that did not pass were excluded from all downstream analyses.

**Figure S4.**
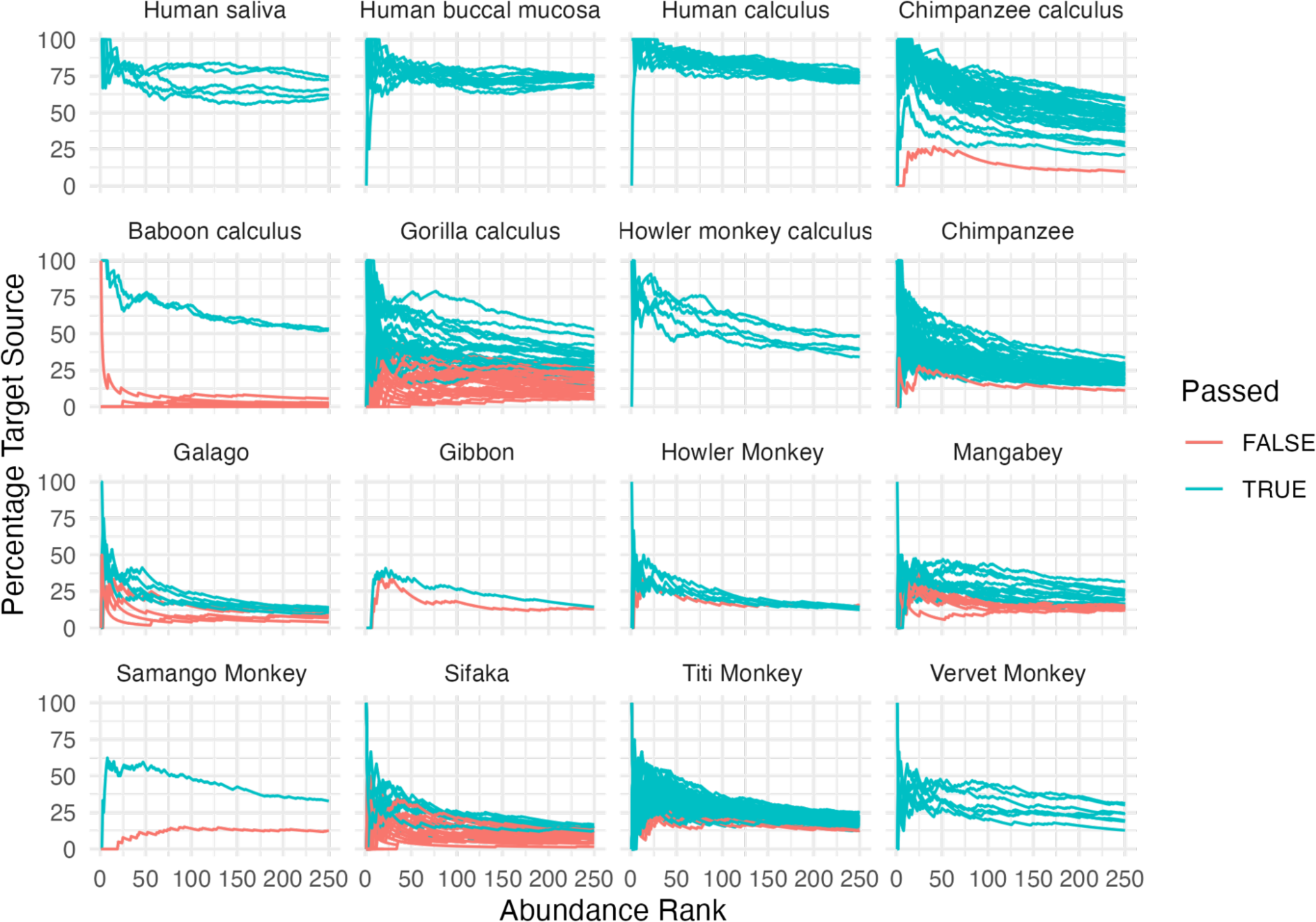
Cumulative percent decay curves for non-human primate historic calculus samples and oral swab samples, with human oral samples for reference. Lines indicate the percent of all species at and above the abundance rank that are classified as oral. Passed - False indicates the sample did not pass the preservation cut-off; Passed - True indicates the sample passed the preservation cut-off and is well-preserved. For consistency with human samples, all non-human primate samples that did not pass were excluded from all downstream analyses. However, the extent of diversity in most non-human primate samples is not known, and these samples may be well-preserved, containing oral species that are not yet characterized, or not yet recognized as oral.

**Figure S5.**
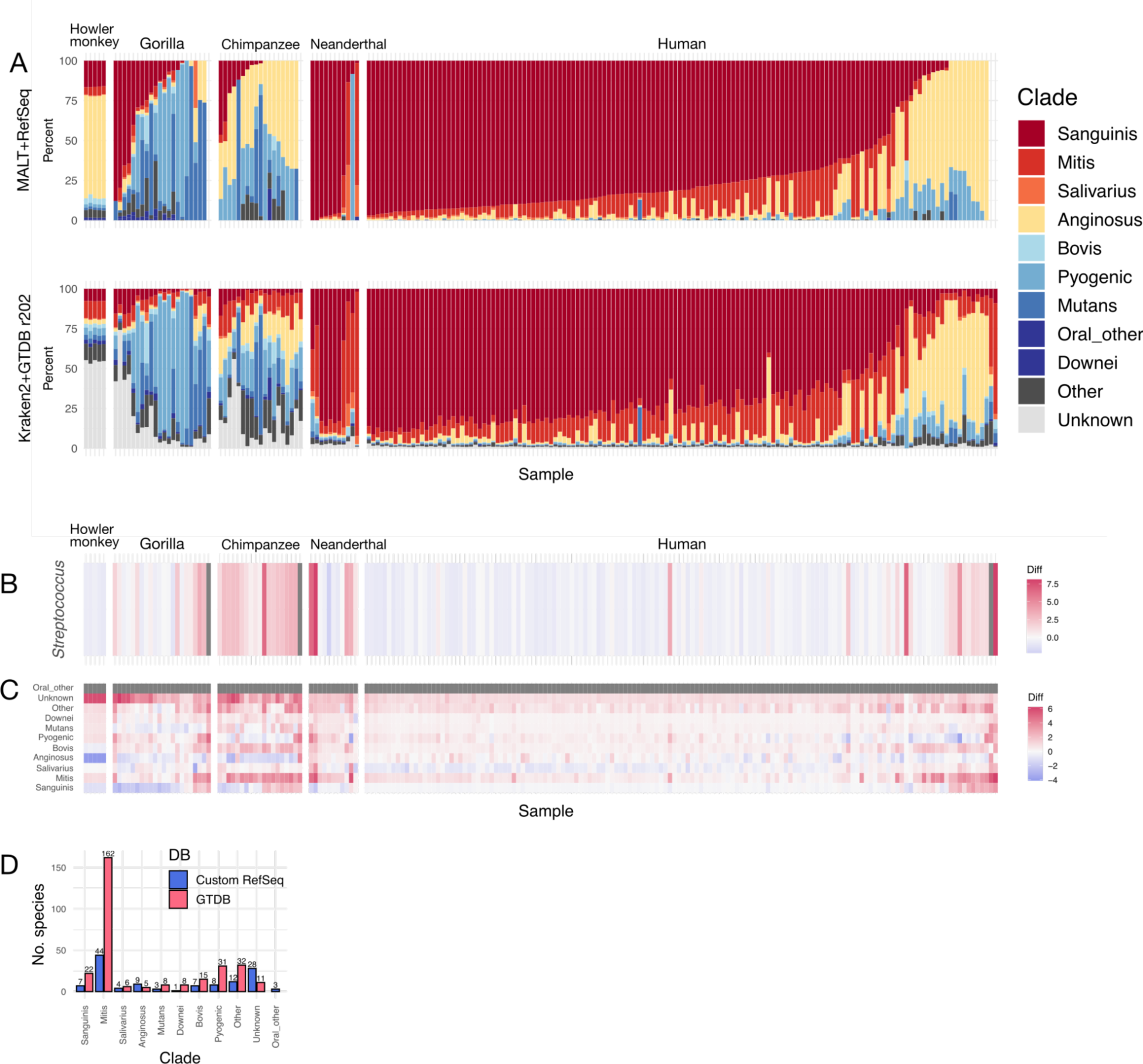
Comparison of *Streptococcus* clades detected using different taxonomic classifiers with different databases on the samples from Fellows Yates, *et al.* 2021^1^. **A.** Top panel - The taxonomic classifier MALT used with a custom NCBI RefSeq database from Fellows Yates, *et al.* 2021. Bottom panel - Kraken2 used with the GTDB r202 database. In the legend, Oral_other refers to the custom RefSeq database only, while Downei refers to the GTDB database only. **B.** Difference in the proportion of reads assigned to species in the genus *Streptococcus* between MALT+RefSeq database and Kraken2+GTDB database for each sample. **C.** Difference in proportion of reads assigned to each Streptococcus clade between MALT+RefSeq database and Kraken2+GTDB databasefor each sample. **D.** Number of species in each *Streptococcus* clade in the custom RefSeq MALT database and the GTDB r202 database.

**Figure S6.**
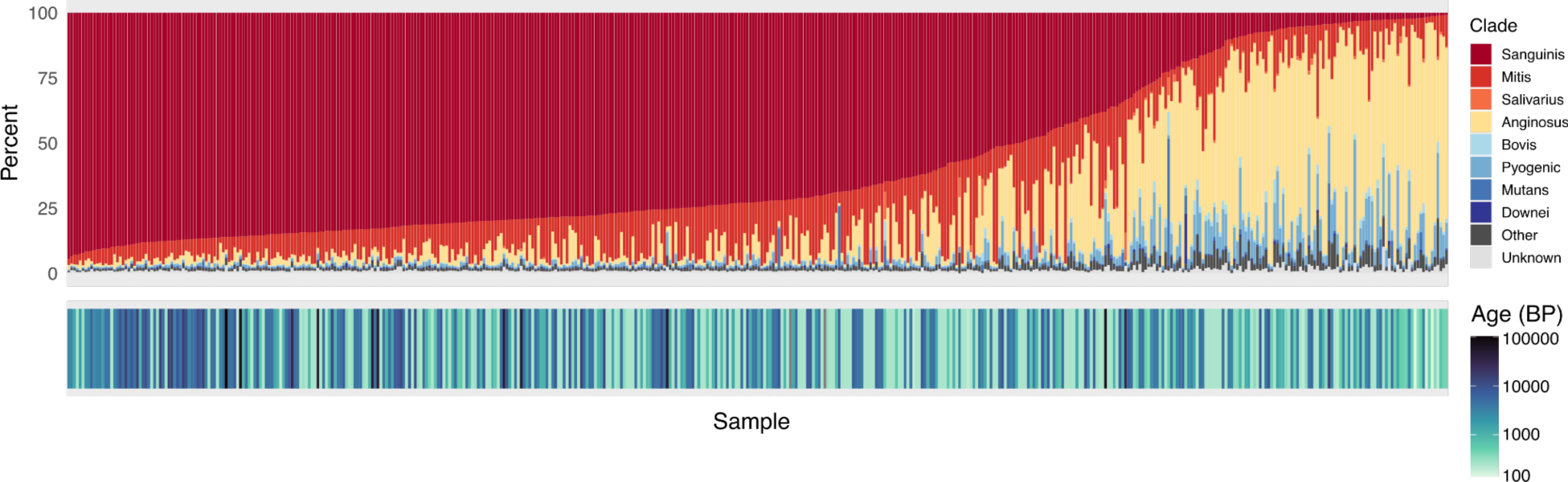
Distribution of *Streptococcus* clades in ancient dental calculus samples with the age of each sample indicated in the lower bar. Sample order is the same as in main text Figure 1.

**Figure S7.**
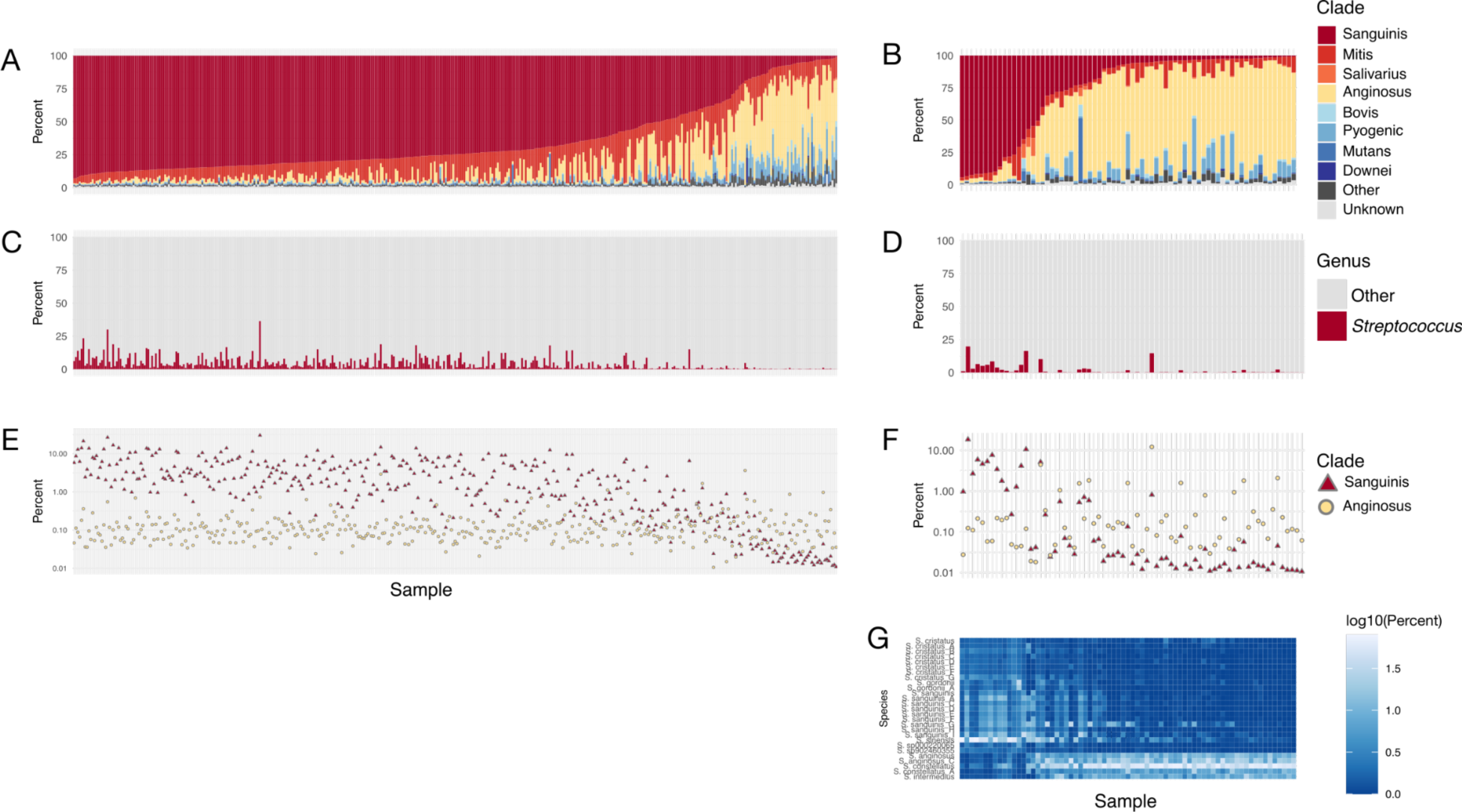
Distribution of *Streptococcus* clades in ancient dental calculus samples. **A, C, E** - all ancient samples except the Pacific calculus dataset from Velsko, *et al.* 2024^10^; **B, D, F, G** - the Pacific calculus dataset from Velsko, et al. 2024. **A,B.** Percent of *Streptococcus* reads that were assigned to each clade, ordered by decreasing abundance of Sanguinis clade and increasing abundance of Anginosus clade. **C,D.** Percent of reads assigned to species in the genus *Streptococcus* and to all other genera. **E,F.** Percent of reads assigned to species in the Sanguinis and Anginosus clades out of al species-level read assignments. **G.** Relative abundance of Sanguinis and Anginosus clade species in the Pacific calculus samples from Velsko, et al. 2024. For Relative abundance of Sanguinis and Anginosus clade species in all ancient calculus samples, see Supplemental Figure S14.

**Figure S8.**
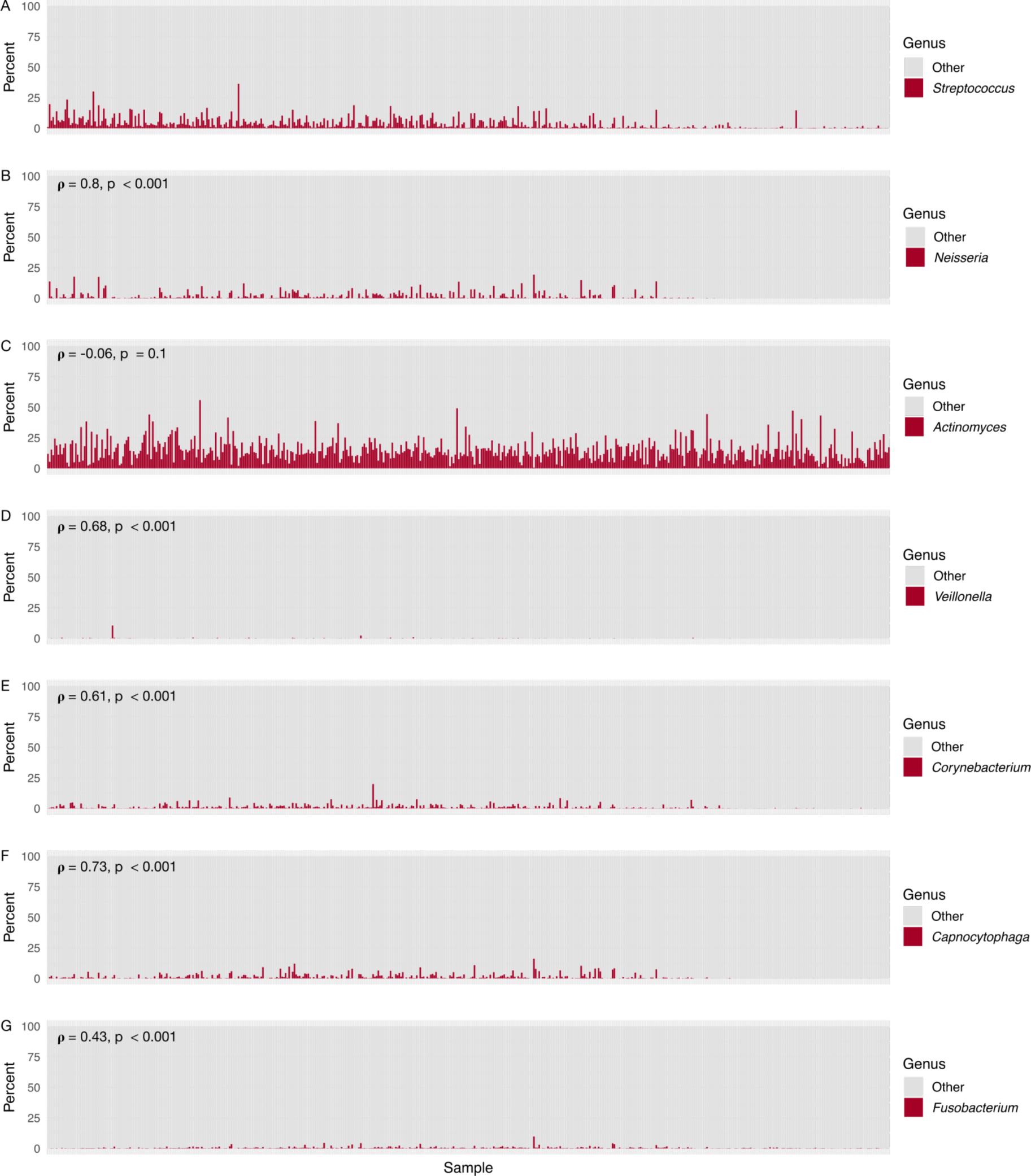
Abundance of genera that include common early colonizer (A-D) and bridging/structural species (E-F) in ancient dental calculus samples. Spearman correlation coefficients (**ρ**) were calculated between the abundances of *Streptococcus* and each other genus, and are indicated in the upper left corner of each plot. Sample order for all plots follows that of main text Figure 1. **A.** Percent of reads assigned to species in the genus *Streptococcus* compared to all other genera, same as main Figure 1B, for reference. **B.** Percent of reads assigned to species in the genus *Neisseria* compared to all other genera. **C**. Percent of reads assigned to species in the genus *Actinomyces* compared to all other genera. **D.** Percent of reads assigned to species in the genus *Veillonella* compared to all other genera. **E.** Percent of reads assigned to species in the genus *Corynebacterium* compared to all other genera. **F.** Percent of reads assigned to species in the genus *Capnocytophaga* compared to all other genera. **G.** Percent of reads assigned to species in the genus *Fusobacterium* compared to all other genera.

**Figure S9.**
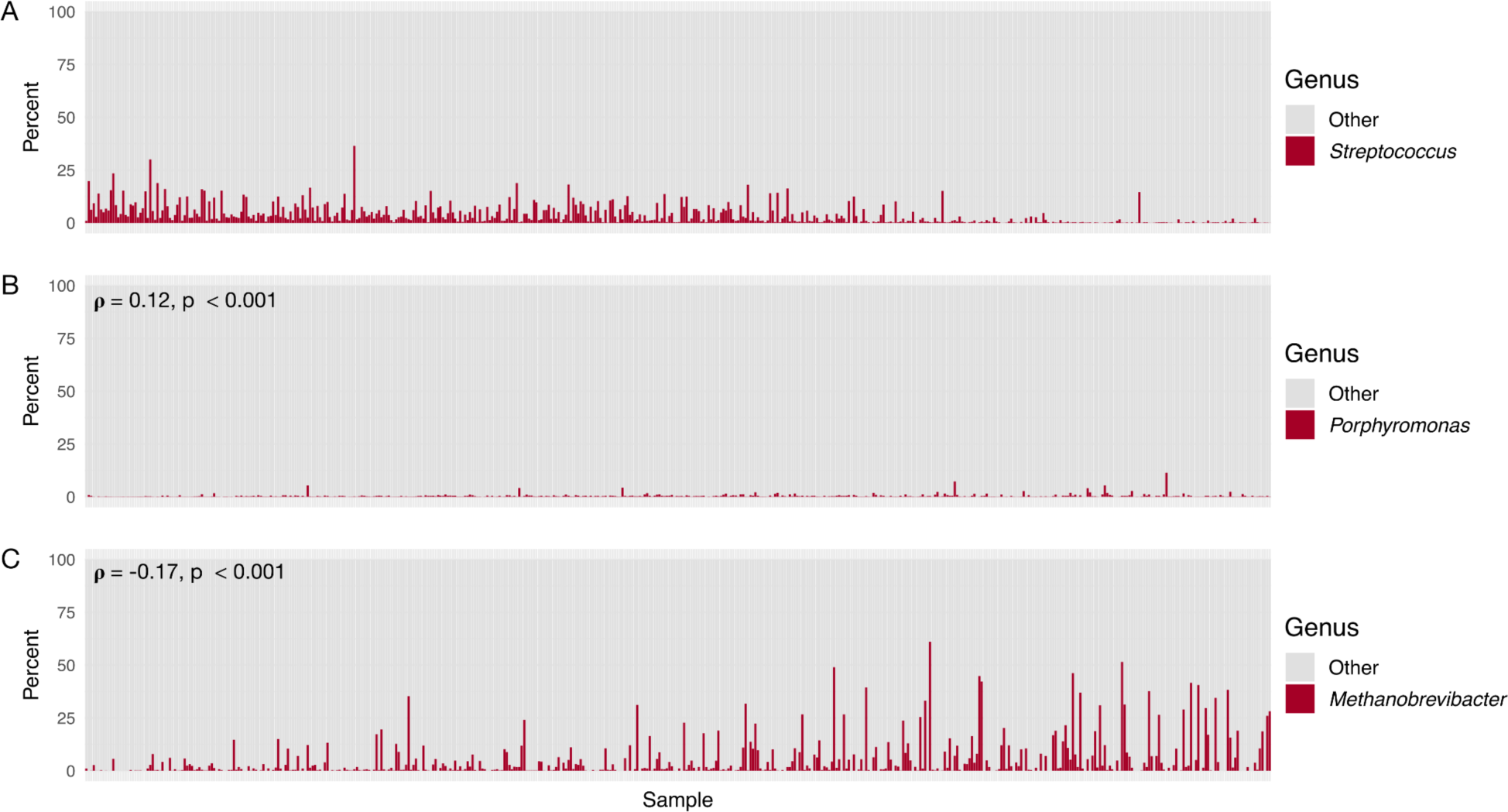
Abundance of late colonizer genera that have described associations with *Streptococcus*. Spearman correlation coefficients (**ρ**) were calculated between the abundance of *Streptococcus* and each other genus, and are indicated in the upper left corner of each plot. Sample order for all plots follows that of main text Figure 1. **A.** Percent of reads assigned to species in the genus *Streptococcus* compared to all other genera, same as main Figure 1B, for reference. **B.** Percent of reads assigned to species in the genus *Porphyromonas* compared to all other genera. A weak positive correlation was detected between the abundance of *Streptococcus* and *Porphyromonas*. **C.** Percent of reads assigned to species in the genus *Methanobrevibacter* compared to all other genera. A weak negative correlation was detected between the abundance of *Streptococcus* and *Methanobrevibacter*.

**Figure S10.**
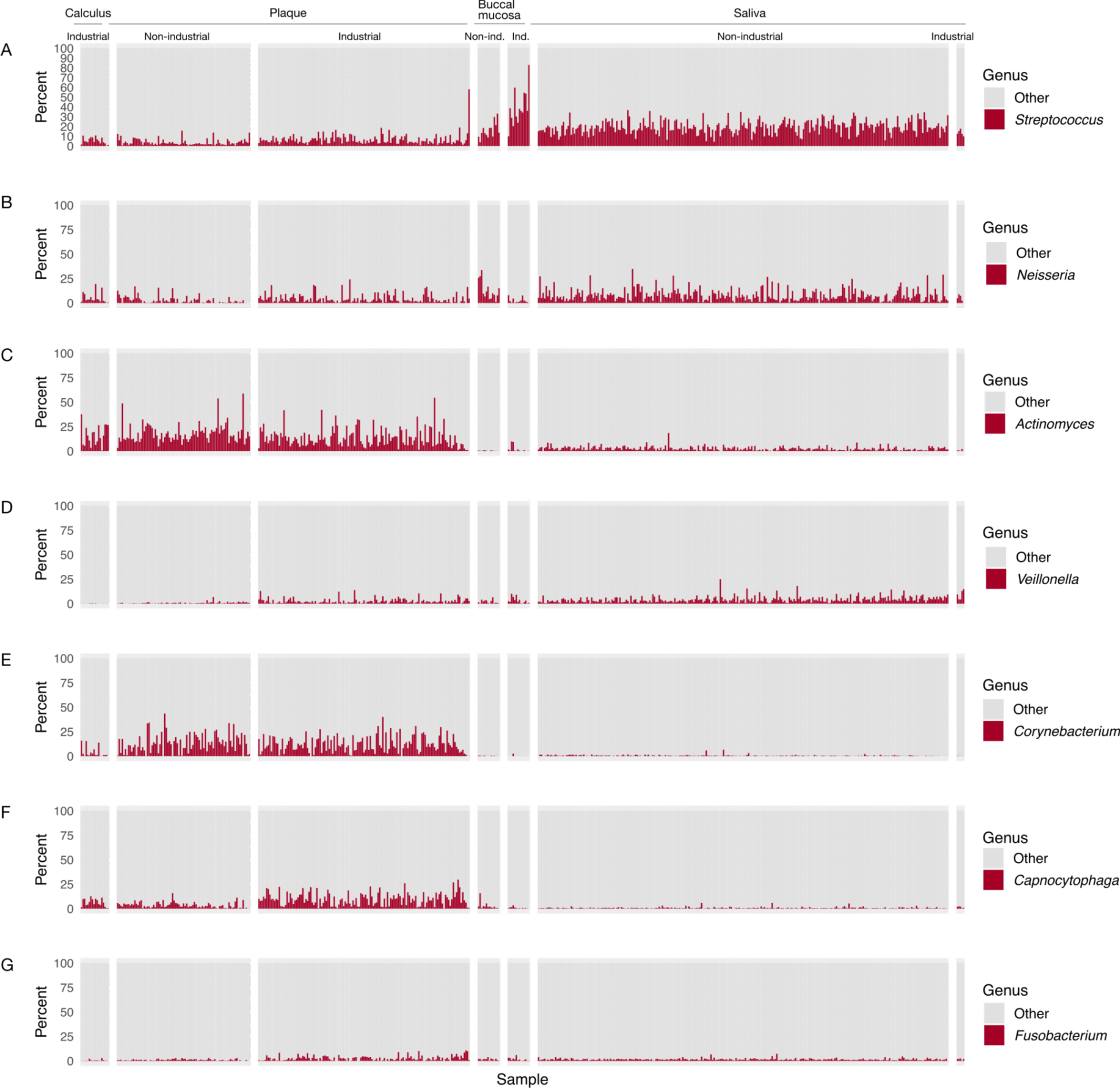
Abundance of genera that include common early colonizer (A-D) and bridging/structural (E-F) species in modern human oral samples. Sample order for all plots follows that of main text Figure 4. **A.** Percent of reads assigned to species in the genus *Streptococus* compared to all other genera, same as main text Figure 4 for reference. **B.** Percent of reads assigned to species in the genus *Neisseria* compared to all other genera. **C**. Percent of reads assigned to species in the genus *Actinomyces* compared to all other genera. **D**. Percent of reads assigned to species in the genus *Veillonella* compared to all other genera. **E**. Percent of reads assigned to species in the genus *Corynebacterium* compared to all other genera. **F**. Percent of reads assigned to species in the genus *Capnocytophaga* compared to all other genera. **G**. Percent of reads assigned to species in the genus *Fusobacterium* compared to all other genera.

**Figure S11.**
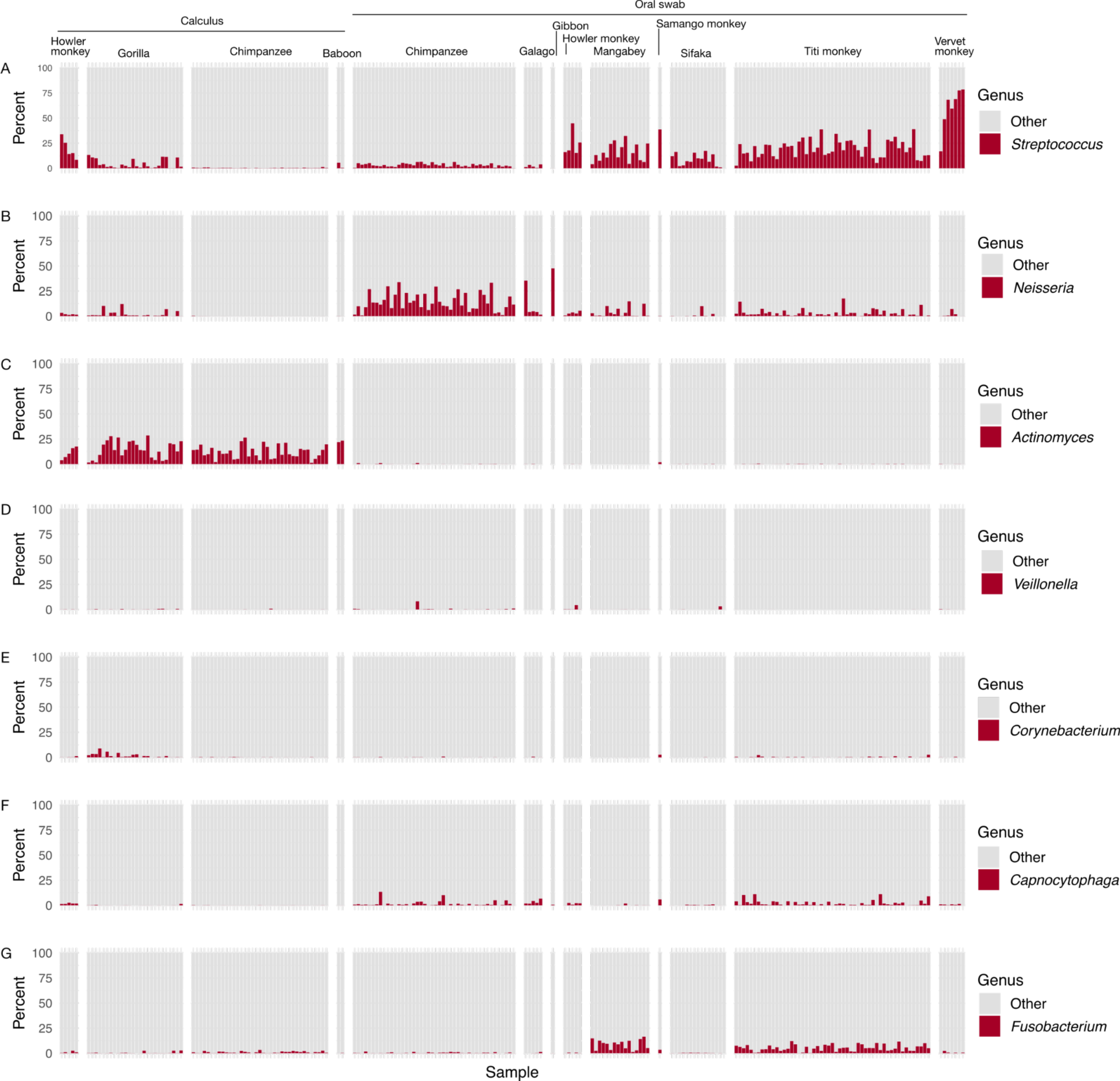
Abundance of genera that include common early colonizer (A-D) and bridging/structural (E-F) species in non-human primate samples. Sample order for all plots follows that of main text Figure 6. **A.** Percent of reads assigned to species in the genus *Streptococcus* compared to all other genera. Same as main text figure 6, for reference. **B.** Percent of reads assigned to species in the genus *Neisseria* compared to all other genera. **C**. Percent of reads assigned to species in the genus *Actinomyces* compared to all other genera. **D**. Percent of reads assigned to species in the genus *Veillonella* compared to all other genera. **E**. Percent of reads assigned to species in the genus *Corynebacterium* compared to all other genera. **F**. Percent of reads assigned to species in the genus *Capnocytophaga* compared to all other genera. **G**. Percent of reads assigned to species in the genus *Fusobacterium* compared to all other genera.

**Figure S12.**
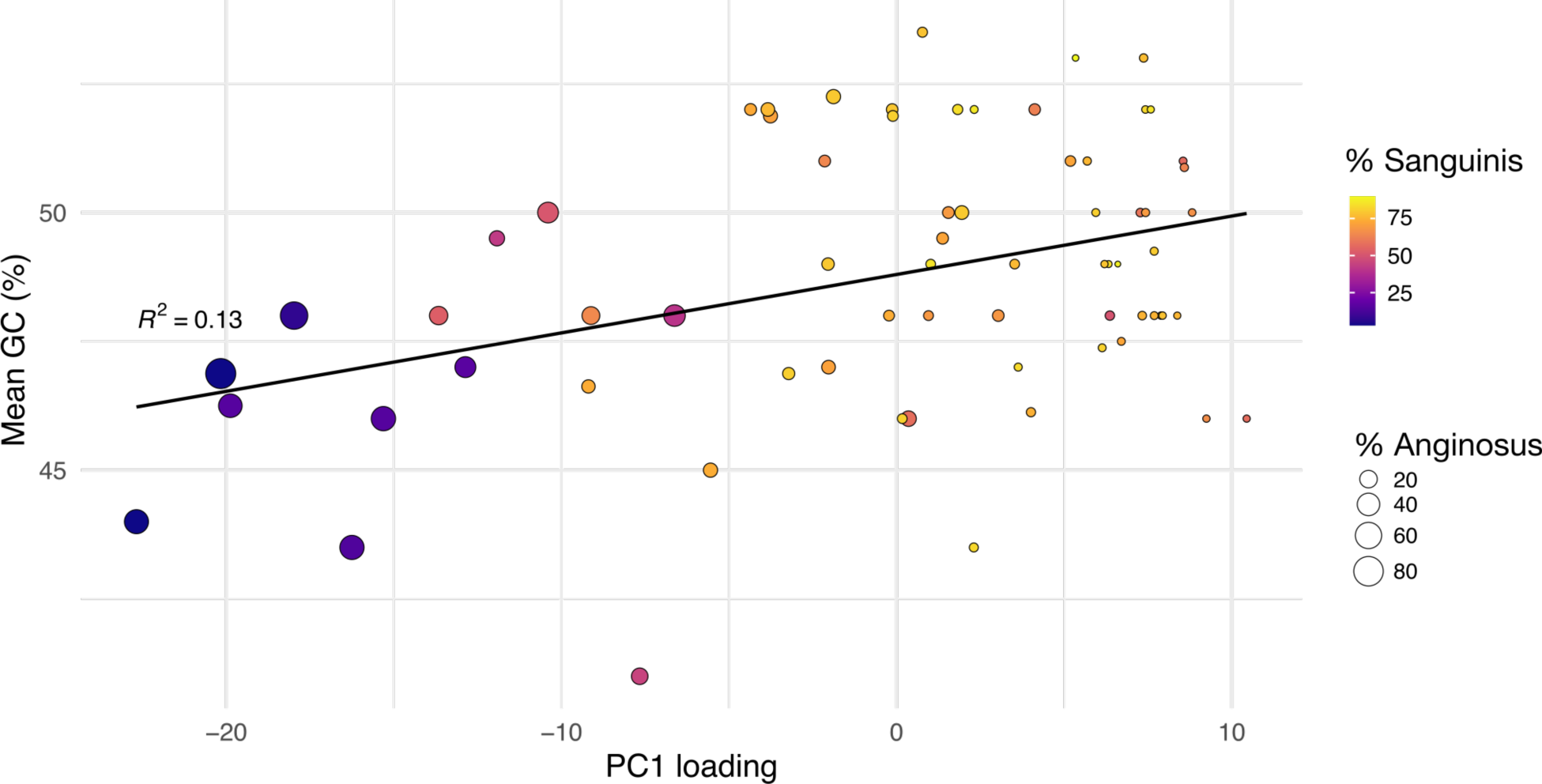
Plot of Middenbeemster samples (featured in main text Figure 2) showing correlations between sample PC1 loadings (x-axis), and the mean sample GC content (y-axis), and the % of reads assigned to the Sanguinis clade (point color), or the Anginosus clade (point size). Linear regression indicates a weakly positive correlation between a sample’s PC1 loading and mean GC content (R^2^ = 0.13). Samples with higher proportions of Sanguinis clade species plot in higher PC1 values and have on average higher GC content than samples with higher proportions of Anginosus clade species, providing a visualization for the correlations reported in main text Figure 2.

**Figure S13.**
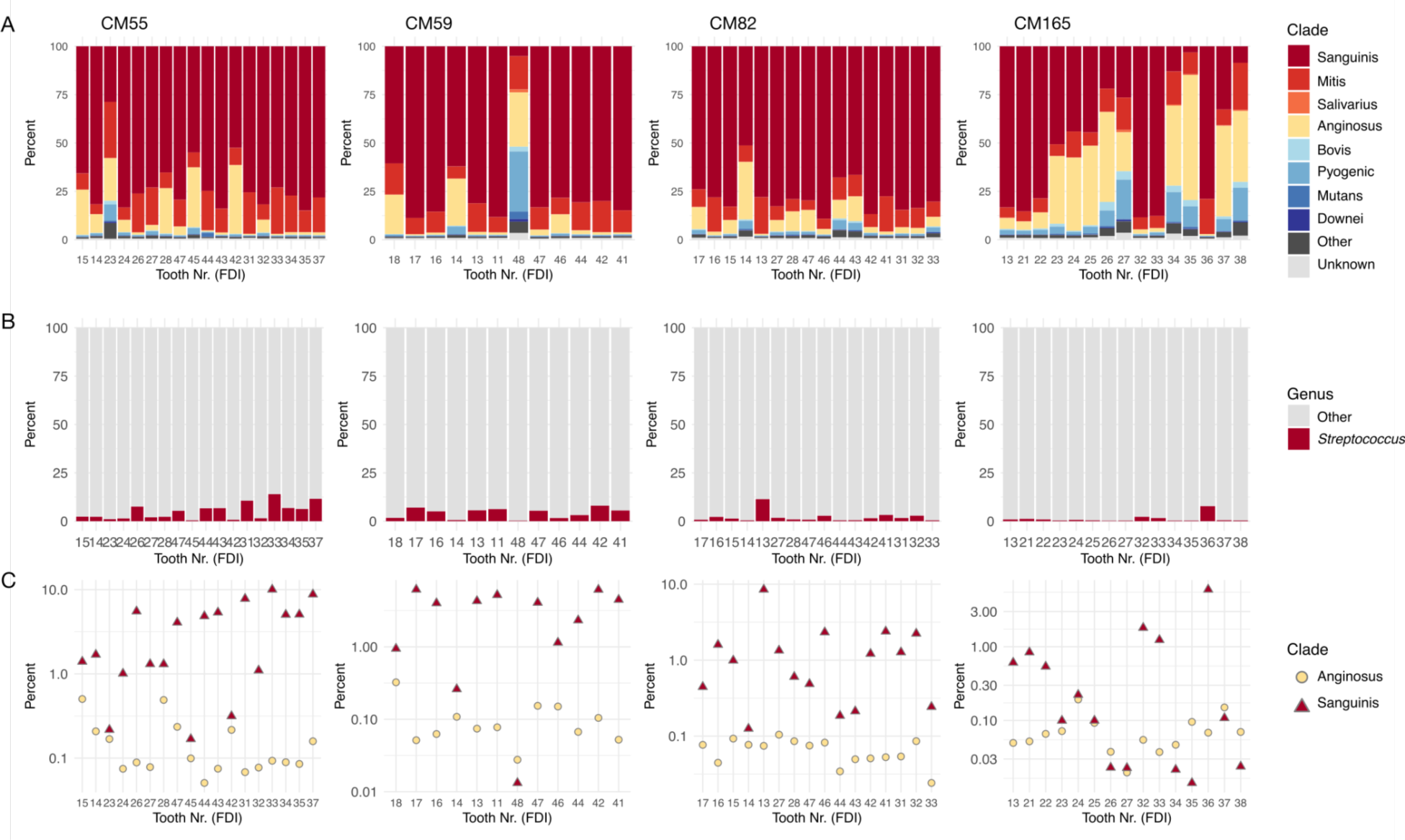
Distribution of *Streptococcus* groups in calculus of each tooth sampled from two individuals from the Chalcolithic site (ca. 4500-5000 BP) Camino del Molino, Spain, expanded for all teeth from all 4 individuals (CM55, CM59, CM82, CM165) used in the original study (Fagernäs, *et al.* 2022^11^). **A.** Percent of *Streptococcus* reads that were assigned to each clade, ordered by decreasing abundance of Sanguinis clade and increasing abundance of Anginosus clade. **B.** Percent of reads assigned to species in the genus *Streptococcus* and to all other genera. **C.** Percent of reads assigned to species in the Sanguinis and Anginosus clades out of all species-level read assignments.

**Figure S14.**
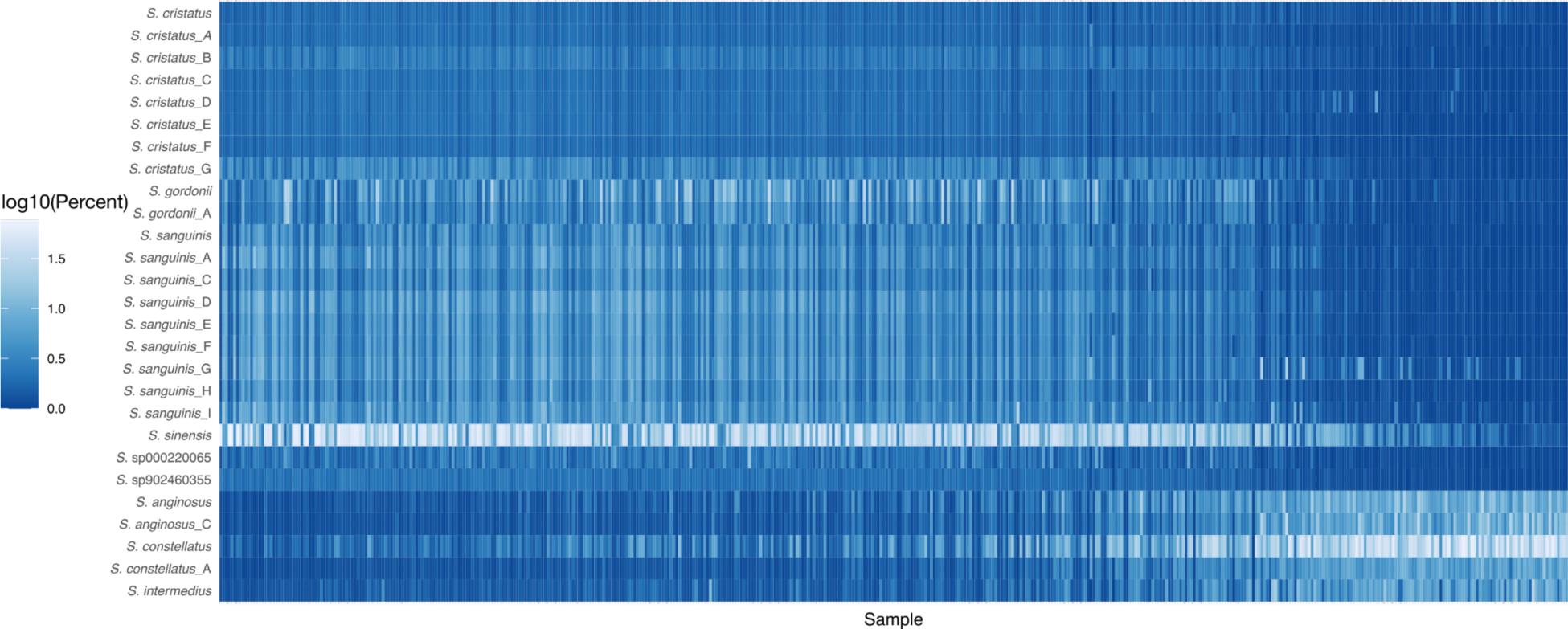
Relative abundance of each of the species of *Streptococcus* in the Sanguinis and Anginosus clades in ancient dental calculus.

**Figure S15.**
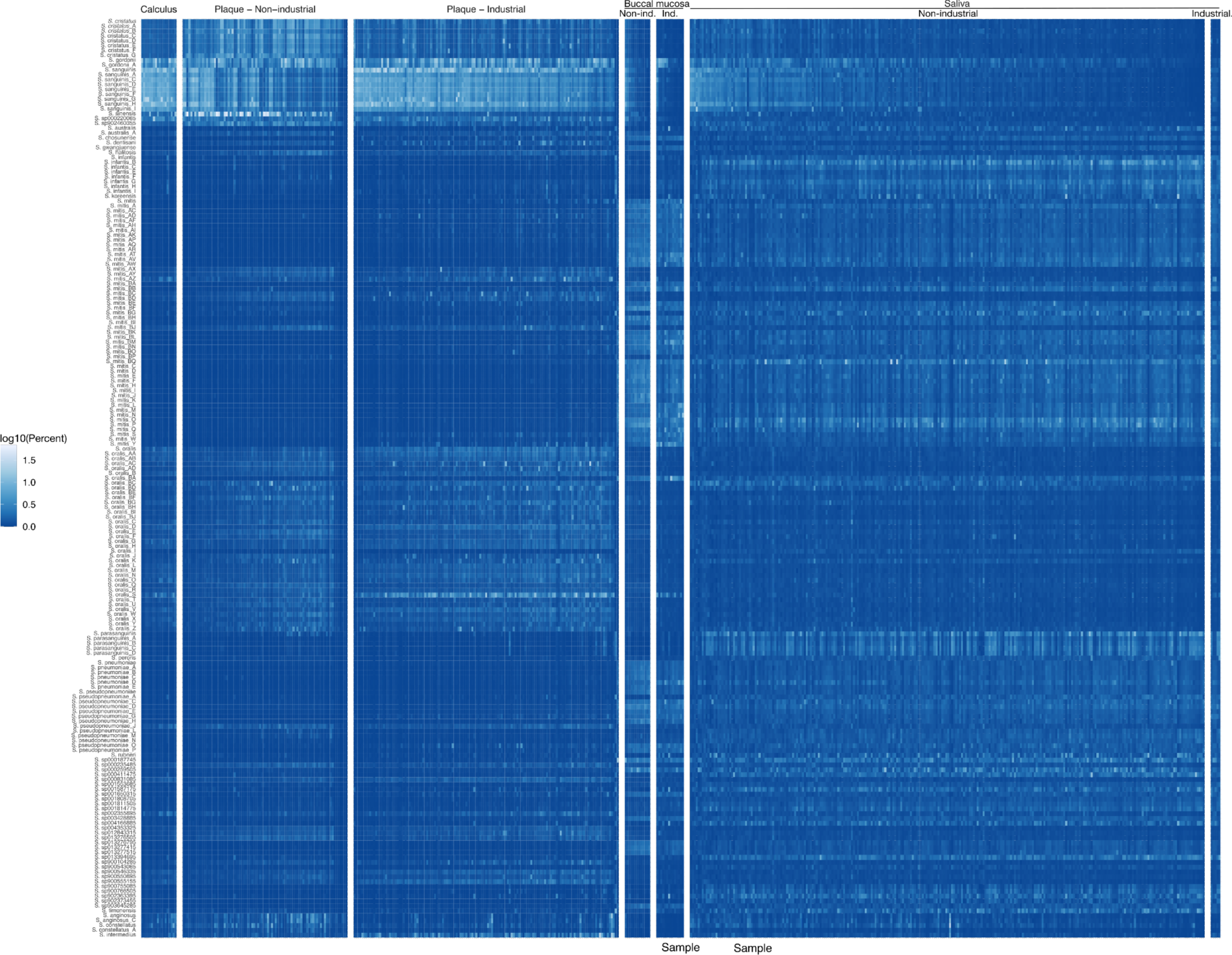
Relative abundance of each of the species of *Streptococcus* in the Sanguinis, Mitis, and Anginosus clades in modern human oral samples. Ind. - Industrial; Non-ind. - Non-industrial.

**Figure S16.**
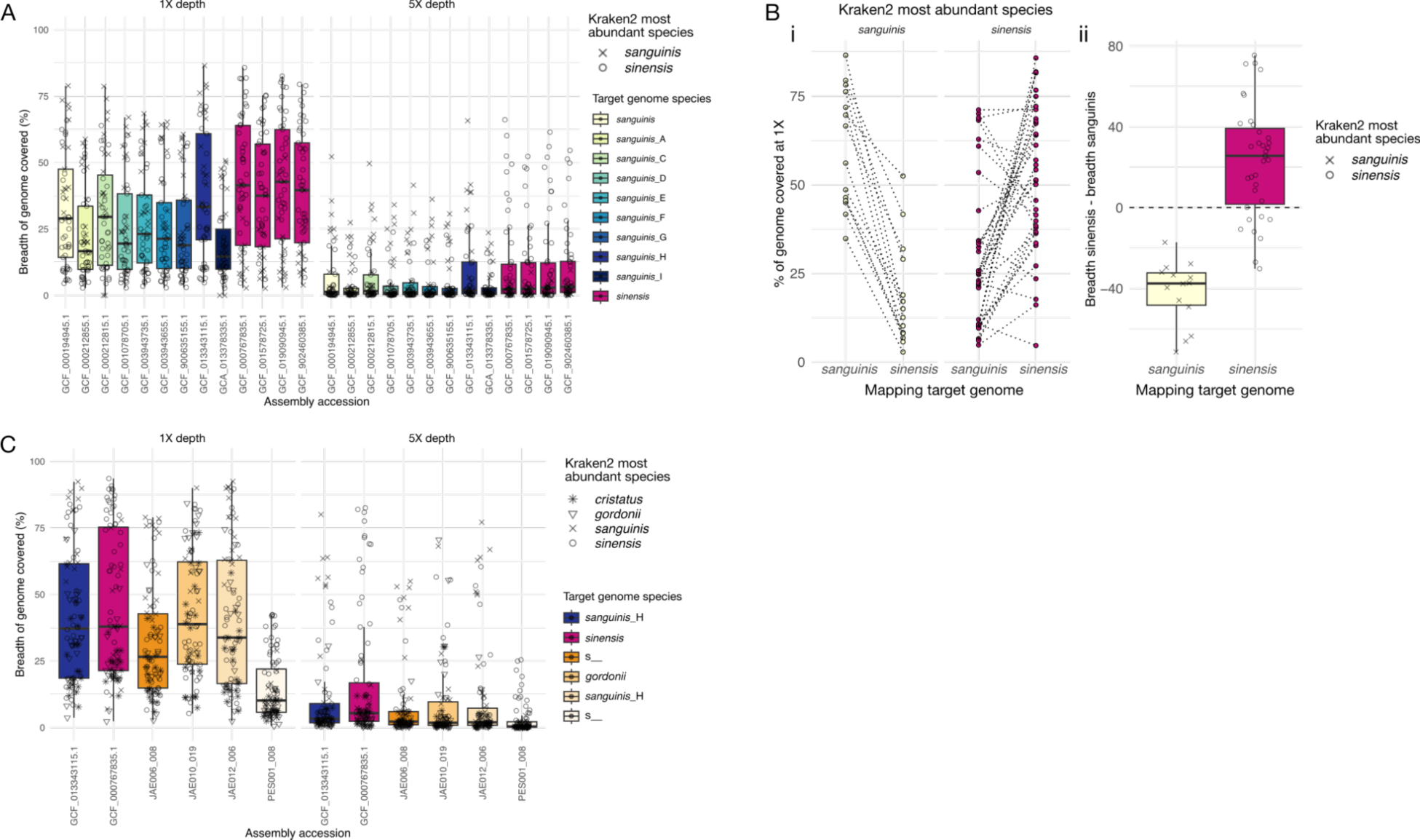
Prevalence and abundance of *S. sanguinis* and *S. sinensis* in ancient dental calculus samples by genome mapping. **A.** Breadth of genome covered at least 1X (left panel) or at least 5X (right panel) by each calculus sample mapped against a concatenated file with representative *S. sanguinis* and *S. sinensis* genomes. Samples are shaped by the species which was most abundant in that sample based on Kraken2 assigned reads. Only samples in which either *S. sanguinis* or *S. sinensis* were the most abundant *Streptococcus* by Kraken2 profiling were plotted. **B. i** - The breadth of target genome (bottom label) covered at least 1X by calculus samples for which *S. sanguinis* (yellow dots) or *S. sinensis* (pink dots) was the most abundant species identified by Kraken2. Samples were mapped against both the *S. sinensis* and *S. sanguinis* genomes, and identical samples are connected with a dotted line. **ii** - the difference in breadth of coverage for each sample in the right panel. All samples for which *S. sanguinis* was the most abundant species by Kraken2 profiling have higher coverage when mapped against a *S. sanguinis* genome than against a *S. sinensis* genome. A majority of samples for which *S. sinensis* was the most abundant species by Kraken2 profiling have higher coverage when mapped against a *S. sinensis* genome than against a *S. sanguinis* genome. **C.** Breadth of genome covered at least 1X (left panel) or at least 5X (right panel) by each calculus sample mapped against a concatenated file of representative *S. sanguinis* and *S. sinensis* genomes as well as four MAGs assembled from modern (JAE) and ancient (PES) dental calculus from Klapper, et al. 2023^7^ that were determined by ANI clustering to belong to the Sanguinis clade. The estimated species for each MAG based on GTDB-TK classification is shown, where s indicates the MAG could not be clustered with a known species in the database. Samples are shaped by the species which was most abundant in that sample based on Kraken2 assigned reads.

**Figure S17.**
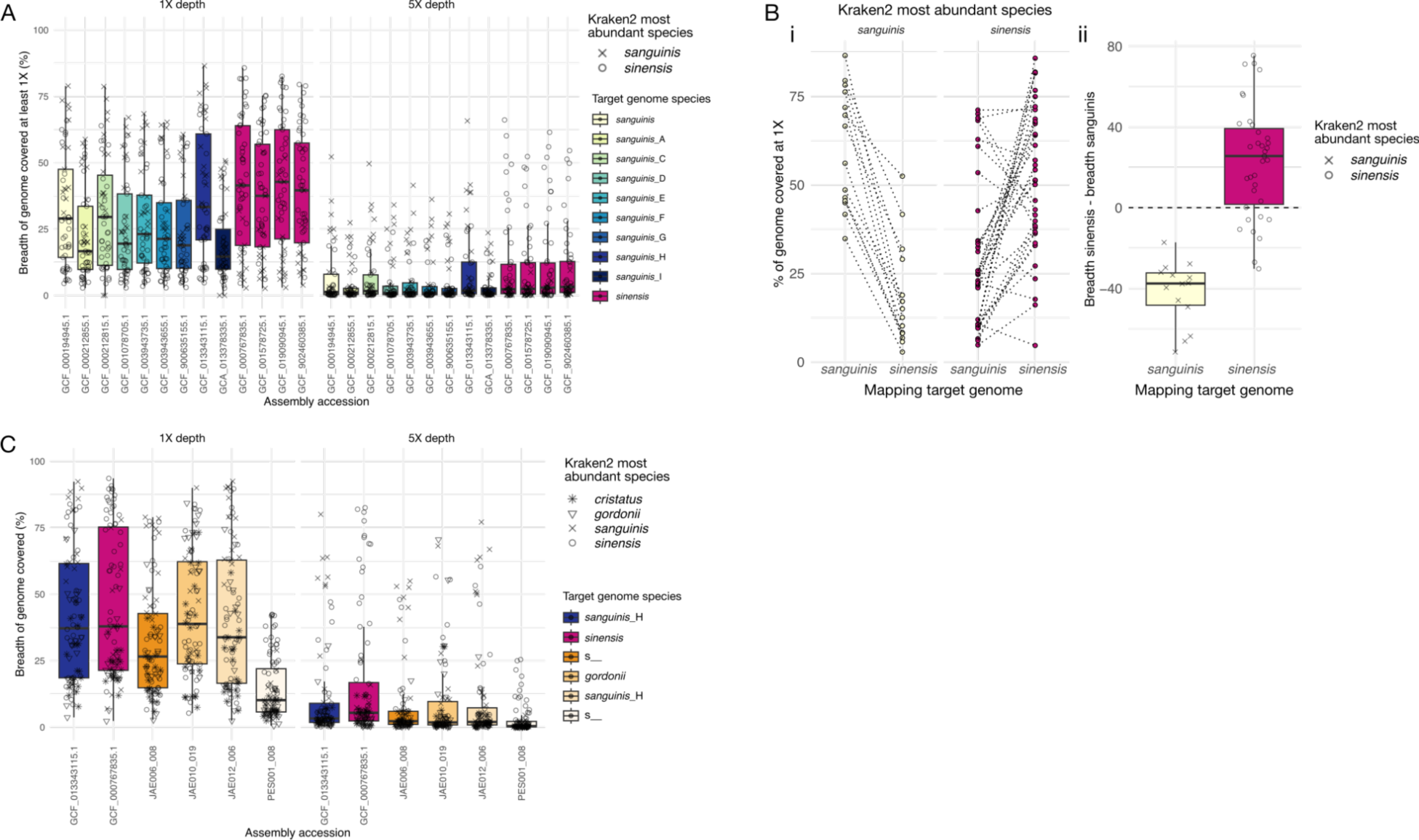
Prevalence and abundance of *S. sanguinis* and *S. sinensis* in Baka and Nzime dental plaque samples by genome mapping. **A.** Breadth of genome covered at least 1X (left panel) or at least 5X (right panel) by each calculus sample mapped against a concatenated file of representative *S. sanguinis* and *S. sinensis* genomes. Samples are shaped by the species which was most abundant in that sample based on Kraken2 assigned reads. Only samples in which either *S. sanguinis* or *S. sinensis* were the most abundant *Streptococcus* by Kraken2 profiling were plotted. **B. i** - The breadth of target genome (bottom label) covered at least 1X by calculus samples for which *S. sanguinis* (yellow dots) or *S. sinensis* (pink dots) was the most abundant species identified by Kraken2. Samples were mapped against both the *S. sinensis* and *S. sanguinis* genomes, and identical samples are connected with a dotted line. **ii** - the difference in breadth of coverage for each sample in the right panel. All samples for which *S. sanguinis* was the most abundant species by Kraken2 profiling have higher coverage when mapped against a *S. sanguinis* genome than against a *S. sinensis* genome. A majority of samples for which *S. sinensis* was the most abundant species by Kraken2 profiling have higher coverage when mapped against a *S. sinensis* genome than against a *S. sanguinis* genome. **C.** Breadth of genome covered at least 1X (left panel) or at least 5X (right panel) by each calculus sample mapped against a concatenated file of representative *S. sanguinis* and *S. sinensis* genomes as well as four MAGs assembled from modern (JAE) and ancient (PES) dental calculus from Klapper, et al. 2023^7^ that were determined by ANI clustering to belong to the Sanguinis clade. The estimated species for each MAG based on GTDB-TK classification is shown, where s indicates the MAG could not be clustered with a known species in the database. Samples are shaped by the species which was most abundant in that sample based on Kraken2 assigned reads.

**Figure S18.**
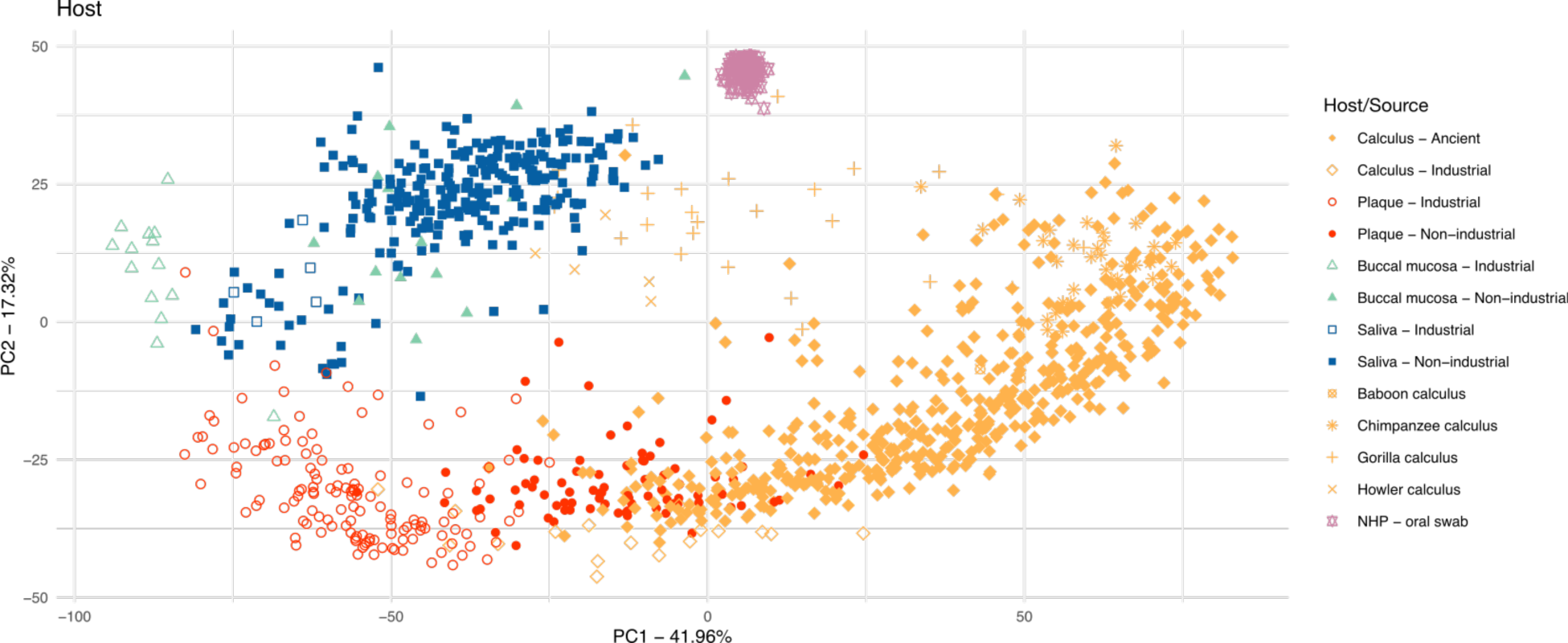
Principal components analysis plot of ancient and modern human and non-human primate oral microbiomes. Shapes and colors indicate sample host. Same plot as main text Figure 7.

